# Distinct patterns of PV and SST GABAergic neuronal activity in the basal forebrain during olfactory-guided behavior in mice

**DOI:** 10.1101/2022.09.04.506524

**Authors:** Elizabeth H. Moss, Evelyne K. Tantry, Elaine Le, Pey-Shyuan Chin, Priscilla Ambrosi, Katie L. Brandel-Ankrapp, Benjamin R. Arenkiel

**Affiliations:** Department of Molecular and Human Genetics, Baylor College of Medicine, Houston TX, 97030, USA; Department of Anesthesiology and Perioperative Medicine, Oregon Health & Science University, Portland, OR, 97239, USA; Neuroscience Graduate Program, Baylor College of Medicine, Houston, TX, 97030, USA

**Keywords:** Olfaction, Basal Forebrain, Parvalbumin, Somatostatin, go/no-go, decision-making

## Abstract

Sensory perception relies on the flexible detection and interpretation of stimuli across variable contexts, conditions, and behavioral states. The basal forebrain is a hub for behavioral state regulation, supplying dense cholinergic and GABAergic projections to various brain regions involved in sensory processing. Of GABAergic neurons in the basal forebrain, parvalbumin (PV) and somatostatin (SST) subtypes serve opposing roles towards regulating behavioral states. To elucidate the role of basal forebrain circuits in sensory-guided behavior, we investigated GABAergic signaling dynamics during odor-guided decision-making. We used fiber photometry to record cell type-specific basal forebrain activity during an odor discrimination task and correlated temporal patterns of PV and SST neuronal activity with olfactory task performance. We found that while both PV-expressing and SST-expressing GABAergic neurons were excited during trial initiation, PV neurons were selectively suppressed by reward whereas SST neurons were excited. Notably, chemogenetic inhibition of BF SST neurons modestly altered decision bias to favor reward-seeking while optogenetic inhibition of BF PV neurons during odor presentations improved discrimination accuracy. Together, these results suggest that the bidirectional activity of GABAergic basal forebrain neuron subtypes distinctly influence perception and decision-making during olfactory guided behavior.

**SIGNIFICANCE STATEMENT:** This study reveals distinct roles for basal forebrain GABAergic neurons in odor perception and odor-guided decision-making. Fiber photometry shows that basal forebrain parvalbumin-expressing neurons are selectively suppressed by rewards, while somatostatin-expressing neurons are activated, establishing the unique recruitment of these GABAergic neurons during behavioral reinforcement. Chemogenetic and optogenetic interventions demonstrate divergent roles for these neuronal subtypes in reward-seeking behavior and odor perception. This research provides new insights into how GABAergic neurons in the basal forebrain shape sensory perception and decision-making.

## INTRODUCTION

Behavioral states potently influence neural circuit function and control information processing from the level of sensory perception to cognition, learning, and memory. The basal forebrain (BF) is involved in behavioral state regulation, with BF circuits controlling directed attention(Sarter and Bruno, 1999; Sarter et al., 2001; Hasselmo and McGaughy, 2004; Herrero et al., 2008), sensory processing (Parikh et al., 2007; Chaudhury et al., 2009; Pinto et al., 2013; Bendahmane et al., 2016; Gritton et al., 2016; Ogg et al., 2018), feeding behavior (Herman et al., 2016; Patel et al., 2019), reward prediction and salience (Lau and Salzman, 2008; Lin and Nicolelis, 2008; Hangya et al., 2015; Ledbetter et al., 2016; Zhang et al., 2019), learning and memory (Muir et al., 1993; Devore et al., 2015; Harrison et al., 2016; Zheng et al., 2018), and arousal (Semba, 1991; Szymusiak, 1995; Sarter and Bruno, 1999; Szymusiak et al., 2000; Lau and Salzman, 2008; Anaclet et al., 2015; Xu et al., 2015). BF projections robustly target both sensory and cognitive circuits and their degeneration is thought to underlie cognitive deficits in disease (Bigl et al., 1982; Grothe et al., 2010; Ballinger et al., 2016; Chaves-Coira et al., 2016; Gielow and Zaborszky, 2017). During learning, BF responses to sensory stimuli are modulated by reward expectation (Lin and Nicolelis, 2008; Hangya et al., 2015; Zhang et al., 2019), supporting that BF circuits integrate sensory input with internal information about reward expectation. Additionally, inhibition of the BF impairs attention in response to motivationally salient stimuli (Tashakori-Sabzevar and Ward, 2018). Intriguingly, both cholinergic and non-cholinergic BF neurons show distinct activity in response to sensory stimuli and play different roles in reward-based learning (Hangya et al., 2015; Patel et al., 2019; Böhm et al., 2020; Nunez-Parra et al., 2020; Hanson et al., 2021). Thus, to better understand how BF circuits mediate state-dependent effects on sensory perception and related behaviors, it is necessary to determine the dynamic activity patterns of specific cell types in the BF and how they are modulated across different behavioral contexts.

The mouse olfactory system is an ideal model for examining the influence of BF modulation on sensory perception and decision-making behavior because both the olfactory bulb and olfactory cortex receive significant cholinergic and GABAergic input from the BF (Woolf et al., 1986; Zaborszky et al., 1986; Nunez-Parra et al., 2013; Sanz Diez et al., 2019; Böhm et al., 2020). Olfactory behaviors, including odor discrimination and odor-guided decision-making can be measured via convenient behavioral assays. Studies that have examined the impact of BF neurons on odor processing have found that inhibiting BF GABAergic neurons impairs fine odor discrimination in a cross-habituation test (Nunez-Parra et al., 2013), and ablating BF cholinergic neurons impairs discrimination in a go/no-go task (Zheng et al., 2022). Moreover, both cholinergic and GABAergic BF neurons respond to odors and reinforcement in go/no-go discrimination tasks (Hangya et al., 2015; Nunez-Parra et al., 2020). We have previously shown that BF GABAergic neuronal activity peaks during odor presentations and is suppressed by reward delivery in a go/no-go task – dynamics which mirror changes in BF cholinergic tone (Hanson et al., 2021). This demonstrates that BF GABAergic neurons are acutely and bidirectionally regulated during odor discrimination tasks with positive reinforcement. These experiments, however, did not differentiate subtypes of GABAergic neurons. Towards understanding how specific BF cell types respond to sensory stimulation, decision-making, and discrimination learning, we used fiber photometry to record BF PV and SST neuronal activity while mice performed an olfactory-cued go/no-go discrimination task. Our data reveal distinct patterns of neuronal activity in PV and SST BF neurons that are uniquely correlated with different aspects of task performance. Further, chemogenetic inhibition of BF SST neurons modestly increased decision bias in favor of reward-seeking, while optogenetic inhibition of BF PV neurons during odor presentations improved odor discrimination. Together these findings highlight the unique contributions of BF PV and SST neurons to odor-guided task performance and decision making.

## MATERIALS AND METHODS

### Mice

Mice were maintained on a 12-h light-dark cycle and were treated in compliance with the ethical guidelines of the US Department of Health and Human Services and approved by IACUC at Baylor College of Medicine and Oregon Health and Science University. Both male and female mice over 2 months of age were included for all experiments. SST-Cre (JAX#: 13044), PV-Cre (JAX#: 17320), and VGAT-Cre (JAX#: 028862) mice were originally purchased from Jackson Laboratories.

### Surgery and viral constructs

Surgical procedures were performed under general anesthesia using isoflurane (4% v/v) induction and maintained under isoflurane with O2 (1-2% v/v) inhalation. Following anesthetic induction, mice were secured to a stereotaxic frame and administered 5 mg/kg of meloxicam prior to surgery. Craniotomies were performed over the horizontal diagonal band of Broca (HDB) at the coordinates from Bregma: ML +/-1 mm; AP 0.7 mm; DV -5.4 mm. For fiber photometry, to target the HDB for viral expression of the calcium-indicator GCaMP8s, cohorts of PV-Cre (N = 3 male and 3 female), SST-Cre (N = 5 male and 3 female), or VGAT-Cre (N = 4 female and 4 male) mice were unilaterally injected with 250 nL of AAV-syn-FLEX-jGCaMP8s-WPRE (addgene plasmid: 162377, serotype 2/9, packaged by the BCM Neuroconnectivity Core). Following viral injection, a custom fiber optic implant (0.48 NA, 200 um core diameter, RWD systems) was placed so that the tip of the fiberoptic was 0.1 mm dorsal to the HDB coordinates used for viral injection. Implants were then fixed with Metabond dental cement (Parkell).

For optogenetic stimulation, PV-Cre mice (N = 5 male and 4 female) were bilaterally injected with 250 nL of AAV-Efla-DIO-ChR2(h134R)::EYFP-WPRE-pA (addgene plasmid: 55640, serotype DJ8, packaged by the BCM Neuroconnectivity Core) at the same coordinates. For optogenetic inhibition, PV-Cre mice (N = 3 males and 4 females) were injected at the same coordinates with 250 nL of AAV-ef1a-Flex-iChloC-2A-dsRed (Addgene plasmid: 70762, serotype DJ8, packaged by the BCM Neuroconnectivity Core). Following viral injection, the same fiber optic implants used for photometry were inserted, bilaterally at 5-degree angles and placed so that the tips of the fiberoptics were 0.3 mm dorsal to the HDB coordinates used for viral injection. Implants were then fixed with Metabond dental cement (Parkell). A subset of mice (N = 2 females) did not receive implants and were used for electrophysiology experiments. AAV(2/9) was used to express GCaMP in PV neurons for photometry experiments, while AAV(DJ8) was used to express opsins in PV neurons for optogenetic experiments. It is possible that different serotypes may have led to different expression levels or targeting of distinct subsets of PV neurons between the photometry and optogenetic manipulation experiments, but the immunofluorescence shown in figures 1 and 6 suggests that this is not the case. Serotypes were consistent across photometry experiments targeting SST and PV neurons and across optogenetic experiments targeting PV neurons, allowing those sets of experiments to be directly comparable.

**Figure 1:**
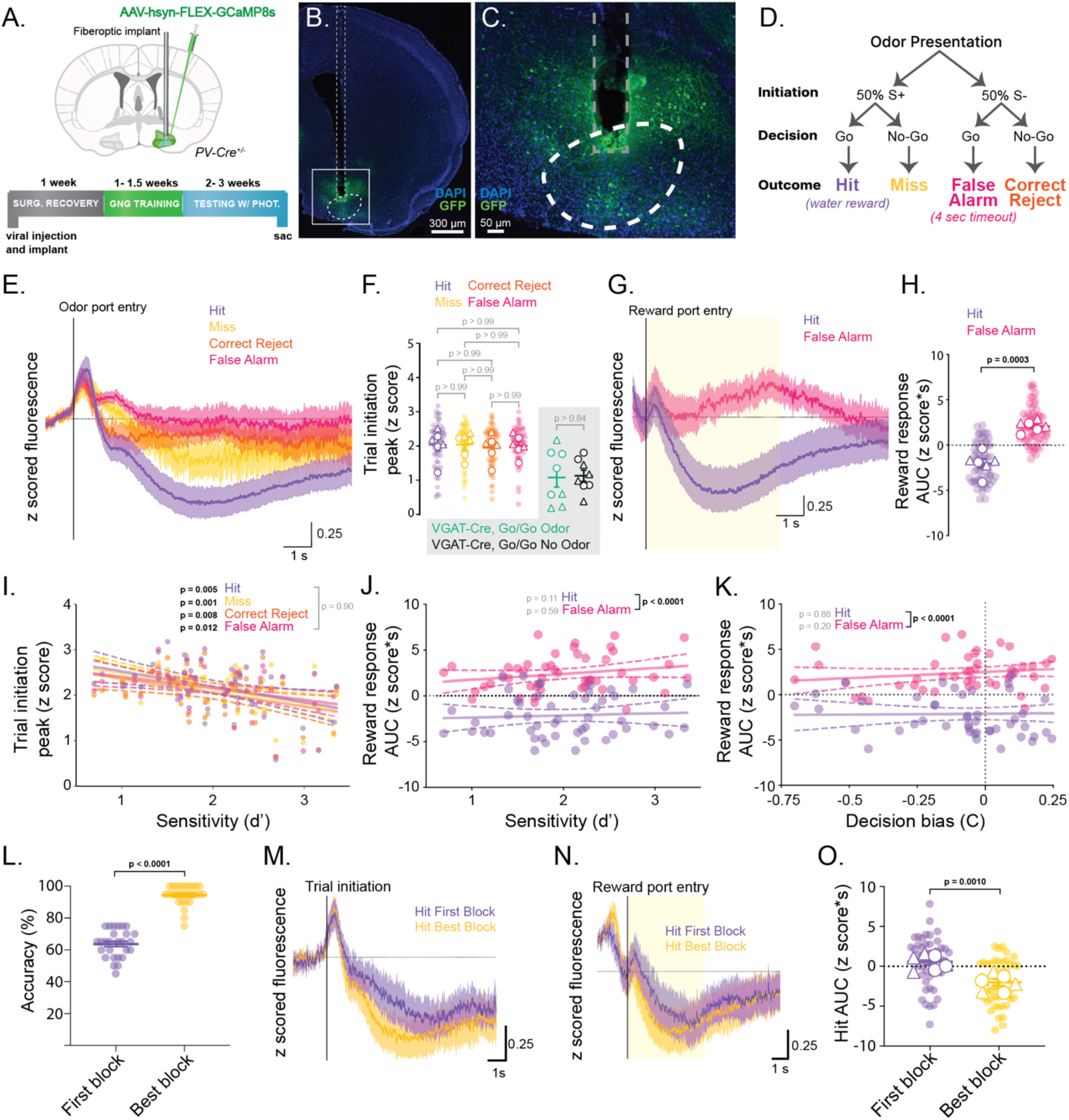
BF PV neurons are excited during trial initiation and suppressed by reward delivery. **A.** Schematic showing injection of Cre-dependent viral construct and fiberoptic implant in PV-Cre mice (top). Highlighted regions of the BF include the horizontal limb of the diagonal band of Broca (HDB), the magnocellular preoptic nucleus (MCPO) and substantia innominata (SI). An experimental timeline shows timing of surgery, behavioral training, and behavioral testing with fiber photometry imaging (bottom). **B.** Example slice image showing the implant tract and expression of GCaMP. **C.** Viral expression (green) and implant tip localization (dashed line) in the HDB. **D.** Schematic of olfactory-cued go/no-go behavioral paradigm showing possible stimuli, decisions, and outcomes of individual trials. **E.** Average z scored GCaMP traces of the four trial types listed in D overlaid with 95% confidence intervals (shaded regions). **F.** (White background) Peak signal amplitudes of the four trial types in the go/no-go task. Pastel circles represent the peak odor response of each testing session. White shapes represent the average peak odor response of each animal. Males are shown as circles and females as triangles. P values comparing go/no-go trial types were calculated using a RM one-way ANOVA with Tukey’s correction for multiple comparisons. Lines show means across animals. Error bars show SEM across animals. (Grey background) Control VGAT-Cre mice performing go/go task with and without odor cues are shown on the right with a grey background for comparison. White markers represent single go/go sessions from each mouse (circles = males, triangles = females). P values were calculated separately from go/no-go trial types using a paired t test. Lines show means across animals. Error bars show SEM across animals. **G.** Average z scored GCaMP traces for Hit and False Alarm trials aligned to the time of reward port entry with 95% confidence intervals overlaid (shaded regions). Yellow shaded area represents the time over which the area under the curve (AUC) was quantified. **H.** Area under the curve (AUC) of Hit and False Alarm traces across testing sessions. Transparent circles represent the AUC of each testing session. White shapes represent the average AUC of each animal with circles representing males and triangles representing females. Lines show mean across animals and error bars show SEM across animals. p = 0.0003 paired t test across N = 6 animals. **I.** Correlation of the peak upon odor port entry with discrimination sensitivity (d’) by trial type, across sessions. Circles show values of odor response peaks for all trials of a given type within a session plotted against the d’ value for that session. Lines of best fit (solid lines) are shown for each trial type with 95% confidence intervals (dashed lines). Slopes of best fit lines that are significantly different from zero are indicated with bolded p values. Grey p value indicates that no best fit line slopes or intercepts are significantly different across trial types. **J.** Correlation of the reward response area under the curve (AUC) with discrimination sensitivity (d’) by reward seeking trial type, across sessions. Circles show values of odor response peaks for hit and false alarm trials within a session plotted against the d’ value for that session. Lines of best fit (solid lines) are shown for each trial type with 95% confidence intervals (dashed lines). Slopes of each best fit line are not significantly different from zero (grey p values). Bracketed, bolded p value indicates a significant difference between the y intercepts of the best fit lines. **K.** Correlation of the reward response AUC with decision bias (C) by reward seeking trial type. Circles show values of odor response peaks for hit and false alarm trials within a session plotted against the C value for that session. Lines of best fit (solid lines) are shown for each trial type with 95% confidence intervals (dashed lines). Slopes of each best fit line are not significantly different from zero (grey p values). Bracketed, bolded p value indicates a significant difference between the y intercepts of the best fit lines. **L.** Accuracy (% correct trials) from first block of 20 trials in a session (purple) compared to the block of 20 trials with highest accuracy in a session (yellow). Circles show values for individual sessions with lines and error bars showing mean and SEM. **M.** Average traces (solid lines), aligned to the time of odor port entry (trial initiation), from hit trials in first blocks (purple) and highest accuracy blocks (yellow) with 95% confidence intervals (shaded regions). Horizontal line indicates baseline subtracted z score = 0. **N.** Average traces (solid lines), aligned to the time of reward port entry, from hit trials in first blocks (purple) and highest accuracy blocks (yellow) with 95% confidence intervals (shaded regions). Horizontal line indicates baseline subtracted z score = 0. **O.** Quantification of the reward-related response (AUC) in hit trials from first (purple) and best blocks (yellow). Circles show values for individual sessions with lines and error bars showing mean and SEM.

For inhibitory DREADD expression, SST-Cre mice were bilaterally injected with 250 nL of AAV-Ef1a-DIO-hM4Di-mCherry (N = 4 males and 4 females, addgene plasmid: 44362, serotype 2/9, packaged by the BCM Neuroconnectivity Core) or AAV-DIO-mCherry (N = 3 males and 3 females, addgene plasmid: 504459, serotype 2/9, packaged by the BCM Neuroconnectivity Core) at the same HDB coordinates. All mice were given a 2-week recovery period prior to the start of behavioral testing. HDB/MCPO/SI targeting was verified in all cases with post hoc immunofluorescence imaging of viral expression.

### Go/no-go training

Freely moving mice were trained on an olfactory-cued go/no-go discrimination task using olfactory conditioning boxes (Med Associates) controlled by MedPC software. Conditioning boxes included one odor delivery port and one water reward port, both equipped with IR beams to record port entries. The odor delivery port received a constant stream of clean room air, cleared by a vacuum. Odors (Sigma, **Tables 1-3**) were injected into the steady air stream by opening valves to one of two odor reservoirs, allowing the air stream to pass through the head space of the reservoir carrying the volatized odorant. Reservoirs were 20 mL in volume and included 2.5 mL of odorant diluted to 1% by volume in mineral oil. Water rewards were dispensed at 5uL in volume in the separate reward port upon reward port entry only on correctly decided Hit trials. Conditioning boxes were housed within larger behavior isolation boxes (Med Associates). Box fans in the isolation boxes continuously circulated ambient air away from the conditioning boxes and out of the isolation boxes to reduce the accumulation of ambient odor.

**Table 1:**
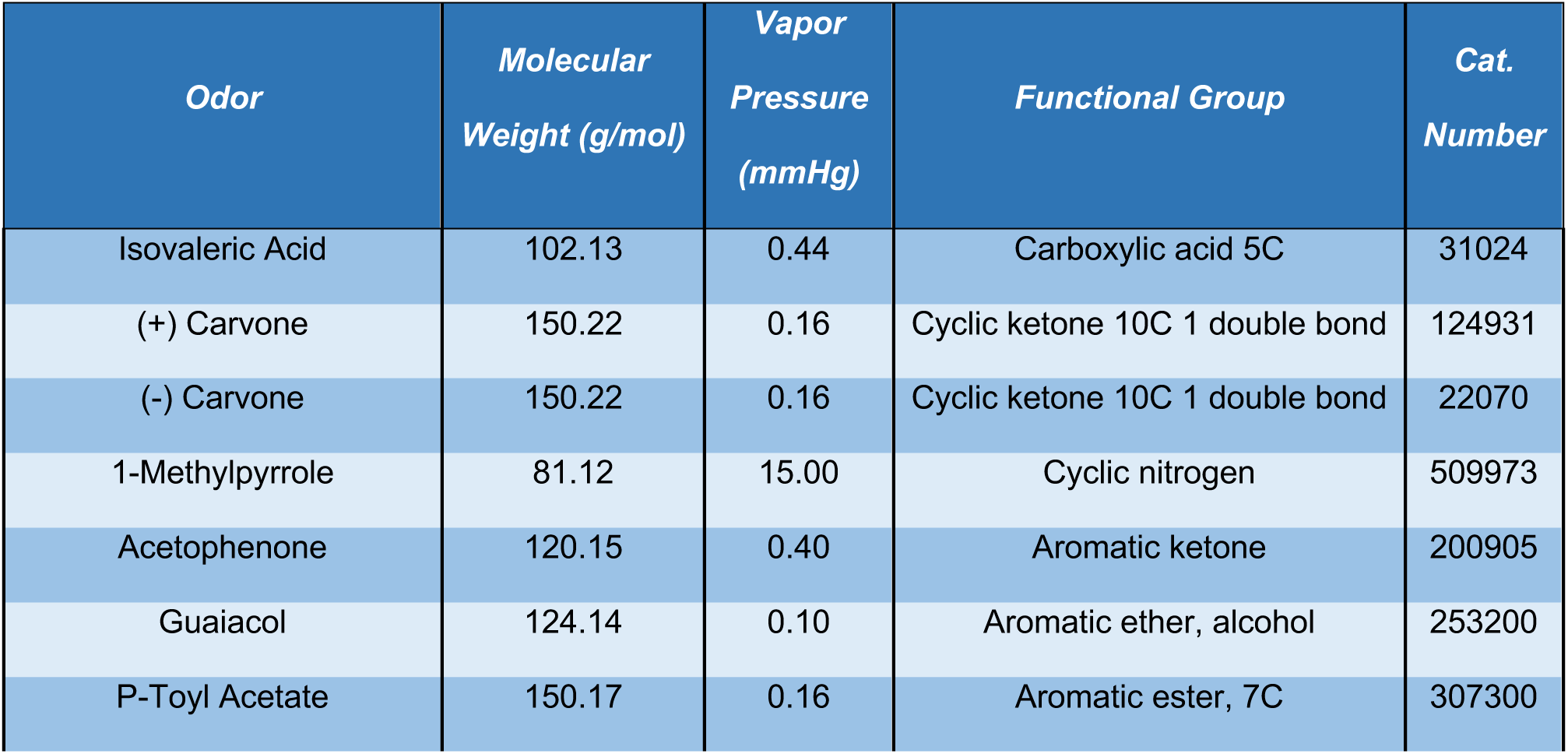
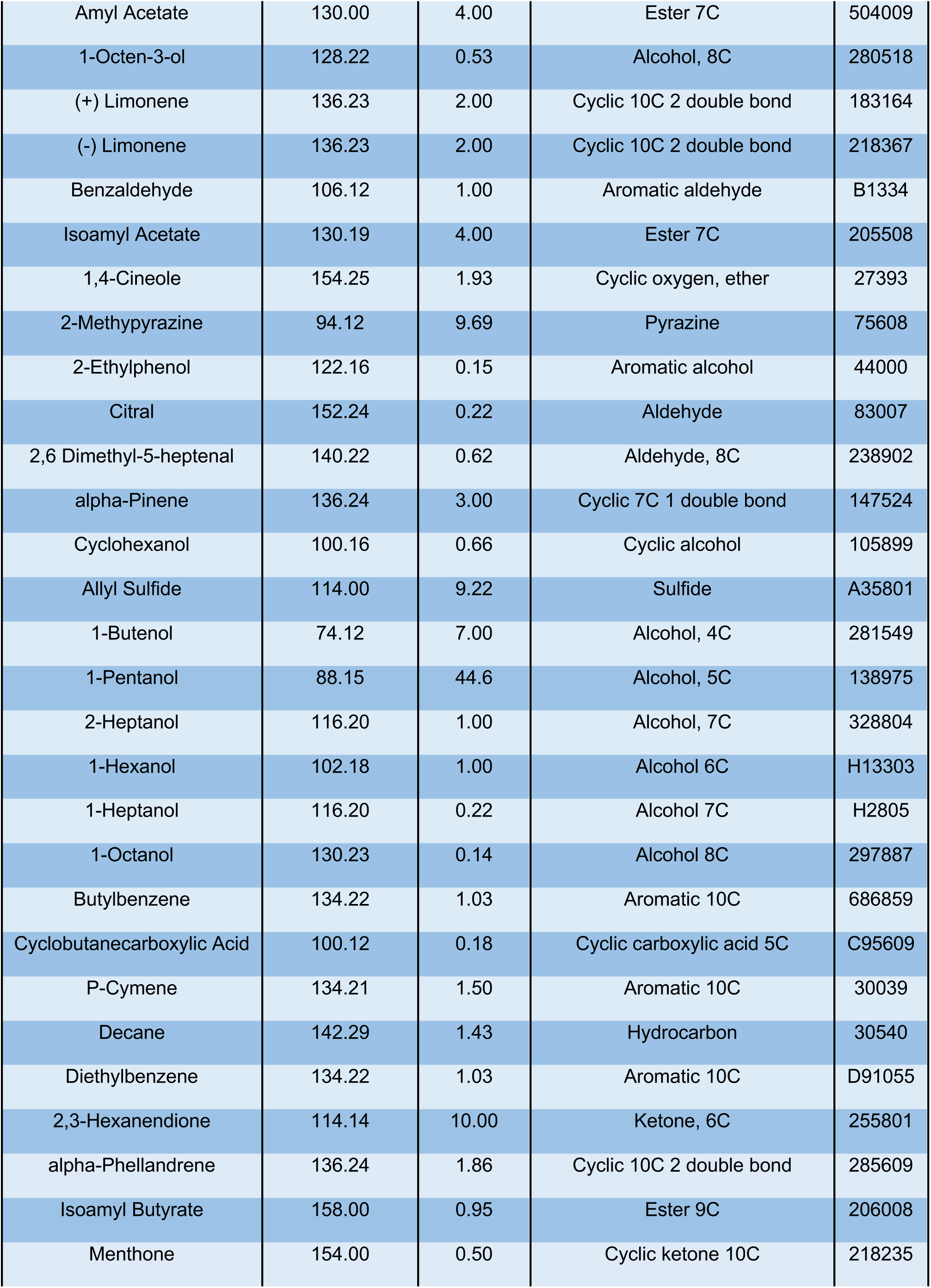
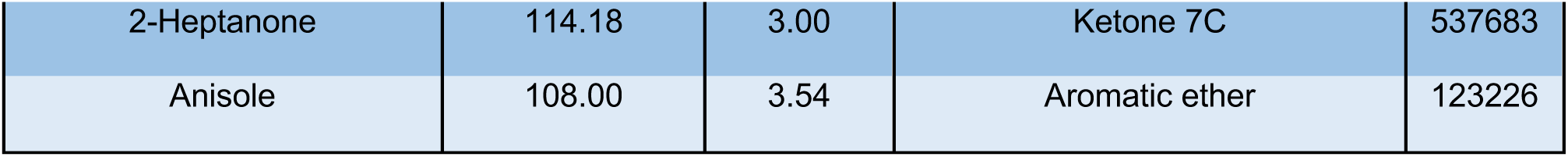
Odors used in go/no-go experiments with fiber photometry.

Over the course of 10-14 days, mice underwent 5 stages of behavior-shaping as previously described (Hanson et al., 2021). Mice were water restricted to no less than 85% of their body weight during the 48 hours prior to the start of training. Mice were first trained to seek water rewards, then to initiate trials by poking their nose into the odor port. Mice were then trained to respond to an S+ odor cue (1% Eugenol in mineral oil) to receive a water reward.

As a control for trial initiation vs. odor detection in photometric recording experiments, a subset of fiberoptic-implanted VGAT-Cre mice underwent photometric recording during a “go/go” version of the behavioral task (**Figure 1F and Figure 2D**). Go/go testing occurred after the training stage where mice had learned to seek water rewards upon odor S+ odor presentations. Instead of continuing with go/no-go training, VGAT-Cre mice underwent single behavioral sessions where in half of the trials they were presented with the S+ odor and in the other half they were presented with no odor. Rewards were available after every trial. In this task, mice obtained rewards with similar high accuracy after both S+ odor and no odor trials.

**Figure 2:**
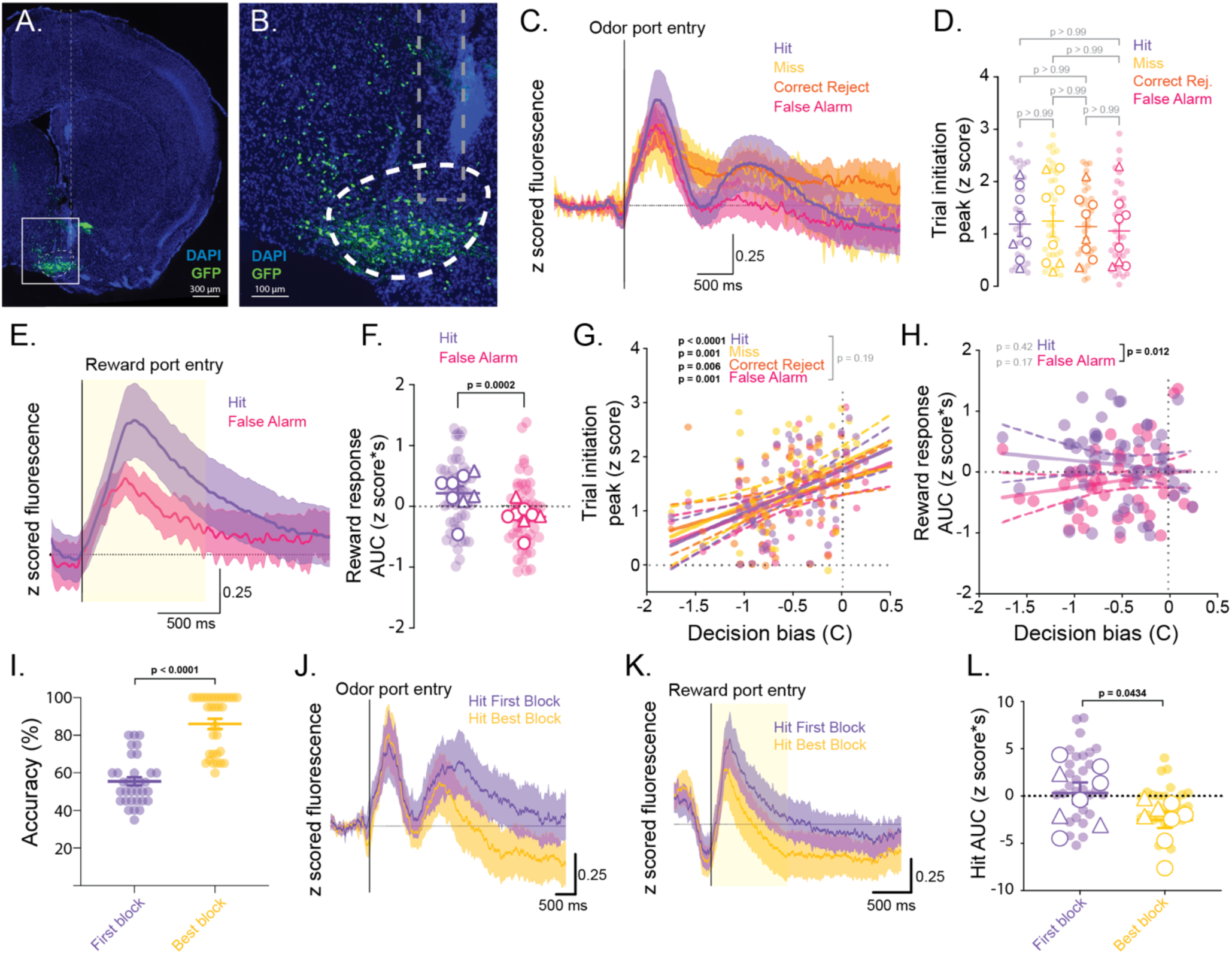
BF SST neurons are excited during trial initiation and reward seeking. **A.** Example slice image showing the implant tract and expression of GCaMP in the BF. **B.** Viral expression (green) and fiberoptic implant tip localization (dashed line) in the HDB. **C.** Average z scored GCaMP traces of the four trial types across all sessions overlaid with 95% confidence intervals (shaded regions). **D.** Peak signal amplitudes of the four trial types upon trial initiation. Light circles represent the peak odor response of each testing session. White shapes represent the average peak response of each animal with mean and SEM of mouse averages. Males are represented as circles and females as triangles. **E.** Average z scored GCaMP traces for Hit and False Alarm trials, aligned to the time of reward port entry, with 95% confidence intervals (shaded regions). Yellow shaded area represents the time over which the area under the curve (AUC) of the reward response was calculated. **F.** Area under the curve (AUC) of Hit and False Alarm traces across testing sessions. Transparent circles represent the AUC of each testing session. White shapes represent the average AUC of each animal (males as circles and females as triangles) with mean and SEM of mouse averages. **G.** Correlation of odor port entry response with decision bias (C) by trial types across sessions. Circles show values of odor response peaks for all trials of a given type within a session plotted against the C value for that session. Lines of best fit (solid lines) are shown for each trial type with 95% confidence intervals (dashed lines). Slopes of best fit lines that are significantly different from zero are indicated with bolded p values. Bracketed, grey p value indicates that no best fit line slopes or intercepts are significantly different across trial types. **H.** Correlation of the reward response AUC with decision bias (C) by reward seeking trial type. Circles show values of odor response peaks for hit and false alarm trials within a session plotted against the C value for that session. Lines of best fit (solid lines) are shown for each trial type with 95% confidence intervals (dashed lines). Slopes of each best fit line are not significantly different from zero (grey p values). Bracketed, bolded p value indicates a significant difference between the y intercepts of the best fit lines. **I.** Accuracy (% correct trials) from first block of 20 trials in a session (purple) compared to the block of 20 trials with highest accuracy in a session (yellow). Circles show values for individual sessions with lines and error bars showing mean and SEM. **J.** Average traces (solid lines), aligned to the time of odor port entry (trial initiation), from hit trials in first blocks (purple) and highest accuracy blocks (yellow) with 95% confidence intervals (shaded regions). Horizontal line indicates baseline subtracted z score = 0. **K.** Average traces (solid lines), aligned to the time of reward port entry, from hit trials in first blocks (purple) and highest accuracy blocks (yellow) with 95% confidence intervals (shaded regions). Horizontal line indicates baseline subtracted z score = 0. **L.** Quantification of the reward-related response (AUC) in hit trials from first (purple) and best blocks (yellow). Circles show values for individual sessions with lines and error bars showing mean and SEM.

For mice undergoing go/no-go training, upon forming an association between the S+ odor and water reward, the S-odor cue (1% Methyl Salicylate in mineral oil) was introduced. In this stage, mice were trained to refrain from seeking a reward and instead initiate another trial. Trials required mice to sample the odor cues for 100ms or more before responding to the cue and were categorized based on the S+/- odor cue and the mouse’s response. When mice were presented with the S+ odor cue, trials were considered “Hits” when mice sought a water reward within 5s and “Misses” if mice did not enter the reward port. Trials in which the S-odor cue was delivered were considered “False Alarms” when mice incorrectly attempted to retrieve water and “Correct Rejects” if mice did not seek water. False alarms resulted in a 4-second time out punishment. Accuracy was calculated as percent correct trials (hit + correct reject trials) for blocks of 20 trials. For each training session, mice completed 10-20 blocks. Training was considered complete after mice completed a session with the training odors (Eugenol = S+, Methyl Salicylate = S-) in which they achieved higher than 85% accuracy in at least 2 consecutive blocks. Testing sessions followed the same pattern as the last stage of training, but new odor pairs were introduced, requiring mice to learn new S+ and S-odor associations (Tables 1-3).

### Go/no-go testing with fiber photometry

For go/no-go testing with concurrent photometry mice were attached to a fiberoptic cable, connected to a two-channel fiber photometry recording system (Doric) via a rotary joint in the top of the behavior box. Odor pairs for photometry experiments were chosen randomly for each mouse and session from the list in Table 1 so that each odor was never presented on more than one day. The timing of trial initiations and reward port entries were measured by IR beams in the odor and reward ports. Port entry timings, odor presentations, and trial outcomes were recorded with (Med Associates). The timing of each odor presentation was output to the photometry recording system as a TTL pulse lasting the duration of an odor presentation allowing synchronization of the photometry and behavior. Sessions were excluded if mice failed to complete at least 100 trials and if the average d’ value across the whole session was less than 0.25.

Fiber optic cables (0.48 NA, 400 um core diameter) connected light emitting diodes (465 nm experimental and 405 nm isosbestic excitation wavelengths) to a filter cube that separating excitation and emission wavelengths (Doric). Excitation wavelengths were directed along a fiber optic tether (0.48 NA, 200 um core diameter) toward the mouse through a rotary connector attached to the behavior box. The same fiber carried emission wavelengths from the mouse to the filter cube, which was then directed to a femtowatt photodetector (Newport) through a fiber optic cable (0.48 NA, 600 um core diameter). 405 and 465 nm excitations were controlled by Doric Studio software in “lock-in” mode and the emission evoked by each channel was separated into distinct Ca^2+^-dependent and isosbestic GCaMP traces in Doric Studio. Traces were then exported to MATLAB where Ca^2+^-dependent GCaMP traces were scaled to the isosbestic traces, isosbestic signals were subtracted to remove artifacts, and the subtracted traces were z scored using the formula *Z* = (*F* − *μ*)/*σ* where *F* = isosbestic-corrected fluorescence, *μ* = the mean of the isosbestic-corrected trace over the entire session, and *σ* = the standard deviation of the isosbestic-corrected trace over the entire session. Signals were then aligned to odor presentations and subdivided into individual go/no-go trials using the TTL output from the behavior apparatus. For display purposes, average traces were baseline subtracted by trial type. Analyses of AUC and peak amplitude were carried out on z scored traces from individual trials without baseline subtraction.

Z scored photometry traces from individual trials within a go/no-go session were sorted based on trial type and outcome. Additional realignment of photometry traces based on reward port entry time was carried out in MATLAB using output from MedPC software. Mean traces were calculated for each session by trial type and averaged across animals and sessions. Error was calculated as the 95% confidence interval across sessions. Peak odor response values were calculated as the maximum value of the z scored photometry trace during the odor presentations on individual trials and then averaged across the session. Session averages were then averaged for each animal and mouse-averaged values were compared across trial types using one way ANOVA. For photometric recordings of GCaMP from SST neurons, the area under the curve (AUC) of the reward-related response was calculated for Hit and False Alarm traces by summing the z score from reward port entry to 1 second after reward port entry. For recordings from PV neurons, reward-related suppression was calculated as the sum of the z scored trace from reward port entry to 5 seconds after reward port entry. Values were not calculated for Miss and Correct Reject trials because those trials did not involve reward port entry. For analyses subdividing sessions to examine within-session learning (**Figure 1L-O**, **Figure 2I-L**), traces were extracted from single blocks of 20 trials. The “first block” included the first 20 trials from a session. The “best block” was considered the block of 20 trials with the highest accuracy. If more than one block achieved the same level of high accuracy, traces from the first instance of high accuracy were used. AUC values were then calculated from Hit trials within the “first block” and the “best block” of a session and averaged by animal. Animal averages were then compared with a paired t-test.

### Go/no-go testing with chemogenetic and optogenetic manipulations

For go/no-go experiments involving DREADD mediated inhibition of SST neurons, odor pairs were presented in the order shown in Table 2. The same odor pairs were presented on consecutive days until mice achieved at least two consecutive blocks of 85% accuracy, when the odor pair was considered learned, or until the mice failed to learn for three consecutive days. Sessions of the same odor pair were then concatenated into a single session spanning up to three days. Concatenated sessions were excluded from analysis if mice failed to complete at least 100 trials. Mice were treated each day, 15 minutes before starting the go/no-go task, with either CNO (3.5 mg/kg in 1% DMSO, Tocris) or vehicle (1% DMSO in saline) by I. P. injection. Mice were habituated to I. P. injections of 1% DMSO during the last stage of go/no-go training. CNO or vehicle treatment was consistent within each concatenated session such that each session of the same odor pair (up to three consecutive sessions) was associated with the same drug treatment (**Figure 4B**). As an additional control, a subset of mice was injected with a non-DREADD expressing virus (AAV-ef1a-FLEX-mCherry) and subjected to the same experimental paradigm.

**Table 2:** Odors used in go/no-go experiments with DREADD-mediated inhibition of SST neurons.

| *Pair #* | *Odor* | *S+/S-* | *Treatment* | *Cat. Number* |
| --- | --- | --- | --- | --- |
| 1 | Anisole | S+ | Vehicle | 123226 |
|  | Acetophenone | S- |  | 200905 |
| 2 | Isoamyl Acetate | S+ | CNO | 205508 |
|  | 2-Heptanone | S- |  | 537683 |
| 3 | 2-Methypyrazine | S+ | Vehicle | 75608 |
|  | alpha-Pinene | S- |  | 147524 |
| 4 | (+) Carvone | S+ | CNO | 124931 |
|  | (-) Carvone | S- |  | 22070 |
| 5 | (+) Limonene | S+ | Vehicle | 183164 |
|  | (-) Limonene | S- |  | 218367 |
| 6 | 2-Ethylphenol | S+ | CNO | 44000 |
|  | 1,4-Cineole | S- |  | 27393 |
| 7 | 1-Butanol | S+ | Vehicle | 281549 |
|  | 1-Pentanol | S- |  | 138975 |
| 8 | 1-Hexanol | S+ | CNO | H13303 |
|  | 1-Heptanol | S- |  | H2805 |
| 9 | 2,3 Hexanendione | S+ | Vehicle | 255801 |
|  | Citral | S- |  | 83007 |
| 10 | Isoamyl Butyrate | S+ | CNO | 206008 |
|  | Menthone | S- |  | 218235 |

For go/no-go testing with optogenetics stimulation of PV neurons, PV-Cre mice injected with conditional ChR2 and implanted with fiberoptics were attached to a bifurcated fiber optic cable attached to commutator at the top of the experimental box that connected to a Doric 473 nm laser. The laser was, in turn, connected to an AMPI Master-8 pulse stimulator that cued the laser to pulse at 20 Hz for 10 ms based on either internal programming or specific behavioral inputs from the mice, depending on the stimulation condition. Laser power was modulated so that the output of the fiber connecting to the fiberoptic implant was 2 mW (∼63 mW/mm^2^ for the 200 □m diameter fiber). Behavioral sessions were run once per day on weekdays. Each session, mice performed the go/no-go behavioral task using one of 12 novel S+/S-odor pairs (**Table 3**). Each session corresponded to one of three different stimulation conditions randomized across sessions and mice so that different mice received different combinations of odor pairs and stimulation conditions. In the “reward-locked stimulation” experimental condition, reward delivery triggered the onset of a 5 second burst of 20 Hz laser light stimulation. In the “regular stimulation” control condition, 5 second pulses of 20 Hz laser light were triggered every 20 seconds regardless of the state of mouse in the behavioral task. In the “no stimulation” control condition, mice were tethered to the bilateral fiberoptic, but no laser light was delivered. Each mouse underwent multiple sessions of all three conditions but was only exposed to each odor pair once.

**Table 3:** Odors used in go/no-go experiments with optogenetic stimulation and inhibition of PV neurons.

| *Pair #* | *Odor* | *S+/S-* | *Cat. Number* |
| --- | --- | --- | --- |
| 1 | Acetophenone | S+ | 200905 |
|  | Anisole | S- | 123226 |
| 2 | Isoamyl Acetate | S+ | 205508 |
|  | Isoamyl Butyrate | S- | 206008 |
| 3 | (+) Carvone | S+ | 124931 |
|  | (-) Carvone | S- | 22070 |
| 4 | (+) Limonene | S+ | 183164 |
|  | (-) Limonene | S- | 218367 |
| 5 | 2-Ethylphenol | S+ | 44000 |
|  | 1,4-Cineole | S- | 27393 |
| 6 | 2-Heptanone | S+ | 537683 |
|  | Menthone | S- | 218235 |
| 7 | 2,3 Hexanendione | S+ | 255801 |
|  | Citral | S- | 83007 |
| 8 | 2-Methypyrazine | S+ | 75608 |
|  | alpha-Pinene | S- | 147524 |
| 9 | 1-Butanol | S+ | 281549 |
|  | 1-Pentanol | S- | 138975 |
| 10 | 1-Hexanol | S+ | H13303 |
|  | 1-Heptanol | S- | H2805 |
| 11 | Guaiacol | S+ | 253200 |
|  | Eucalyptol | S- | 246506 |
| 12 | Trans-cinnamaldehyde | S+ | 239968 |
|  | Methyl Acetate | S- | 296996 |

For go/no-go testing with optogenetics inhibition of PV neurons, PV-Cre mice were injected with conditional iChloC and implanted with fiberoptics then attached to a bifurcated fiber optic cable and commutator that connected to a Doric 473 nm laser as for optogenetic stimulation. For iChloC experiments, an AMPI Master-8 pulse stimulator cued the laser to output a single 500 ms duration pulse, based on either internal programming or specific behavioral inputs from the mice, depending on the stimulation condition. Laser pulse parameters were chosen to maximize inhibition of PV neurons during odor presentations and were informed by prior studies demonstrating lasting inhibition from a single light pulse (Wietek et al., 2015). We also verified effective, lasting inhibition of BF PV neurons in brain slices with electrophysiology (**Figure 5A-D**). Behavioral sessions were run once per day on weekdays. Each session, mice performed the go/no-go behavior task using one of 12 novel S+/S-odor pairs (**Table 3**), similar to optogenetic stimulation experiments. Each session corresponded to one of three different stimulation conditions randomized across sessions and mice so that different mice received different combinations of odor pairs and stimulation conditions. In the “all trials” experimental condition, odor presentations triggered the onset of the 500 ms pulse of laser light. In the “random trials” odor presentations triggered laser pulses 50% of the time at random. In the control condition, 500 ms pulses of laser light were delivered every 20 seconds regardless of the state of the mouse in the behavioral task. Each mouse underwent multiple sessions of the three conditions but was only exposed to each odor pair once.

### Quantification of go/no-go task performance

Go/no-go task performance was quantified using the signal detection theory measures d prime (*d’* = *Z*(*Hit Rate*) – *Z*(*False Alaram Rate*)) and criterion (*C* = -1 (*Z*(*Hit Rate*) + *Z*(*False Alaram Rate*))) Go/no-go task performance was quantified using the signal detection theory (Stanislaw and Todorov, 1999). d’ measures the sensitivity of discrimination and reflects overall task performance. C measures discrimination bias. More negative C values reflect a bias towards reward seeking, values close to 0 reflect optimal decision bias, and positive values reflect bias away from reward-seeking, in favor of trial reinitiation. To show change in sensitivity over time within a session, d’ was calculated over a rolling window of 40 trials. Session summary d’ values were calculated as the maximum d’ over a 40-trial rolling window. This was done to avoid early trials where mice were learning new S+ and S-associations and later trials where mice may have disengaged with the task. For experimental and control mice in SST DREADD inhibition experiments, d’ and C were calculated for each odor pair and values were averaged for each mouse separated by treatment condition. Mean values for each mouse from CNO treatment sessions were compared to mean values from vehicle treatment sessions by paired t-tests. For optogenetic stimulation and inhibition experiments, d’ and C were calculated for each odor pair and values were averaged for each mouse separated by stimulation conditions. Stimulation conditions were then compared using repeated measures one-way ANOVA and Tukey corrections for multiple comparisons.

### Electrophysiology

For slice electrophysiological recording experiments testing DREADD-mediated inhibition of SST neurons in the basal forebrain, SST-Cre mice (N = 3 males) were bilaterally injected with 250 nL of AAV-Ef1a-DIO-hM4Di-mCherry. After two-three weeks, mice were deeply anesthetized with isoflurane then transcardially perfused with ice-cold artificial cerebrospinal fluid (aCSF) solution containing (in mM): 125 NaCl, 2.5 KCl, 1.25 NaH2P04, 1 MgCl2, 2 CaCl2, 25 glucose, and 25 bicarbonate (pH 7.3, 295 mOsM). Brains were removed and transferred into ice-cold cutting solution containing (in mM): 2.5 KCl, 1.25 NaH2PO4, 10 MgSO4, 0.5 CaCl2, 234 sucrose, 11 glucose, and 26 bicarbonate. Cutting solution was continuously bubbled with 95% CO2 / 5% O2. Brains were mounted with cyanoacrylate glue and immediately submerged in oxygenated cutting solution on a Leica VT1200 vibratome. 300 um coronal sections were made at a cutting speed of 0.1 mm/s. Slices including the HDB were removed to a slice recovery chamber of oxygenated aCSF at 37C for at least 30 minutes. Following recovery, slices were slowly returned to room temperature for 30 minutes before recording.

For whole-cell current clamp recordings, slices were submerged in a recording chamber and continuously perfused with room temperature oxygenated aCSF at ∼2 mL/minute. Cells were visualized with DIC optics (Olympus BX50WI) and fluorescence imaging (Q Capture). Once visualized, cells were whole-cell patched in current-clamp configuration. Recording electrodes (3–7 megaohms) were pulled from thin-walled borosilicate glass capillaries (inner diameter: 1.1 mm, outer diameter: 1.5 mm) with a horizontal micropipette puller (Sutter Instruments). Current-clamp internal solution contained (in mM): 120 K gluconate, 20 HEPES, 10 EGTA, 2 MgATP, and 0.2 NaGTP (with 0.4% biocytin by weight, pH to 7.3 with KOH, 285 mOsM). Recordings were made using PClamp software (Axon) with an Axon MultiClamp 700B amplifier digitized at 10 kHz (Axon Digidata 1440A). Once a whole cell patch was achieved, cells were allowed to equilibrate for five minutes before resting membrane potential was recorded and current injections were performed. Cells were then injected with 1 s current steps from -100 pA to +180 pA in intervals of 20 pA. Resting membrane potential was recorded and current injections were repeated three times. After recording current injections at baseline, CNO (10 uM) was washed added to the bath and allowed to circulate for 10 minutes. Resting membrane potential was recorded and current injections were repeated three times in the presence of CNO. During all recordings, access resistance was continuously monitored, and cells where the access resistance changed by more than 20% were excluded from analysis. N = 5 fluorescently labeled cells and N = 7 unlabeled cells. Traces were then exported to MATLAB where resting membrane potential and spike counts were calculated with custom scripts.

For slice experiments testing iChloC-mediated inhibition of PV neurons in the basal forebrain, PV-Cre mice (N = 2 females) were bilaterally injected with 250 nL of AAV-ef1a-Flex-iChloC-2A-dsRed (e^12^ gc/ml). After three weeks, mice were deeply anesthetized with isoflurane and transcardially perfused with ice-cold N-Methyl-D-Glucamine (NMDG) aCSF containing (in mM): 92 NMDG, 2.5 KCl, 1.25 NaH2PO4, 30 NaHCO3, 20 HEPES, 25 Glucose, 5 Na-Ascorbate, 2 Thiourea, 3 Na-Pyruvate, 10 MgSO4, and 0.5 CaCl2 (Thermo Scientific). All extracellular solutions were saturated with 95%O_2_/5%CO_2_ and their pH and osmolarity were adjusted to 7.3-7.4 and 300±5 mOsm, respectively. Brains were quickly removed, glued (Krazy Glue) to the specimen tube of a Compresstome (Precisionary VF-510-0Z) and covered in 1.8% low-melting point agarose (diluted in PBS, kept at 42°C). Agarose was quickly solidified using a chilling block. Coronal slices (300 μm) were cut in NMDG aCSF at speed 2 (4 mm/s) and oscillation 4 (18 Hz vibration frequency) with a stainless-steel blade (Electron Microscopy Science). Slices were allowed to recover for 45 min in three 15 min baths: (1) warm (33°C) NMDG aCSF; (2) warm (33°C) recovery aCSF, containing (in mM): 92 NaCl, 2.5 KCl, 1.25 NaH2PO4, 30 NaHCO3, 20 HEPES, 25 Glucose, 5 Na-Ascorbate, 2 Thiourea, 3 Na-Pyruvate, 2 MgSO4, 2 CaCl2; and (3) room temperature recovery aCSF. Finally, slices were kept at room temperature in recording aCSF, containing (in mM): 125 NaCl, 25 NaHCO3, 1.25 NaH2PO4, 2.5 KCl, 1 MgCl2, 2 CaCl2, 12.5 Glucose. During recordings, aCSF was recirculated at ∼4 ml/min (Gilson Minipuls 2) and warmed to 30-32°C with an inline heater (Warner Instruments). Cell attached and whole-cell current clamp recordings were done with an internal solution containing (in mM): 135 KMeSO3, 10 HEPES, 5 KCl, 2.5 NaCl, 0.5 EGTA, 2 Mg-ATP, 0.3 Na-GTP, 5 Na2-phosphocreatine (pH 7.3-7.4, 290 ±5 mOsm, with 0.4% biocytin). Patch pipettes (3-6 MΩ) were pulled (Sutter P-1000) from borosilicate glass (World Precision Instruments 1B150F-4) and moved with the assistance of a micromanipulator (Sutter). Cells were visualized with a 20x water-immersion objective (NA 1, Olympus #N2699600) on a microscope (Olympus BX51WI) equipped with infrared-differential interference imaging (DIC) and a camera (Excelitas pco.panda 4.2). An LED light source (CoolLED pE-300^ultra^) was used to illuminate the slice through the objective for targeted patching and for optogenetic stimulation. With the aid of a power meter (Thor Labs PM100D and S170C), the LED power was adjusted to deliver ∼60 mW/mm^2^ at 460 nm to the slice during optogenetic stimulation. Recordings were made at 20 kHz in PClamp (Axon) with a MultiClamp 700B amplifier and Digidata 1440A digitizer (Molecular Devices). For whole cell recordings, pipette capacitance neutralization and bridge balance were adjusted. Data analysis was performed offline using custom-written MATLAB scripts (N = 5 fluorescently labeled cells and N = 3 unlabeled cells).

All cells were dialyzed with 0.4% biocytin for the duration of the recording and patched neurons were saved for post hoc imaging. After recordings, electrodes were withdrawn slowly allowing the cells to reseal and form an outside-out patch. Slices were then allowed to equilibrate in the recording chamber for 5 minutes before being transferred to 4% PFA. After patching and cell-filling, slices were fixed overnight in 4% PFA at 4° C. Slices were then washed 3x in 0.1% PBS-T for 30 min each. After washing, slices were incubated in 10% normal goat serum blocking buffer for 2 hours at room temperature. Slices were then incubated in streptavidin conjugated to Alexa:647 (1:500, Invitrogen) overnight at 4C. The next day, slices were washed 3x in 0.1% PBS-T for 30 min, then mounted on glass slides using 500 um spacers (Electron Microscopy Sciences) filled with mounting media without DAPI (Southern Biotech). Slices were imaged on a Leica SP8 confocal with a 20x objective to verify colocalization with DREADD expression. Z stacks of filled cells were reconstructed in FIJI.

### Immunofluorescence and imaging

For antibody staining of GCaMP in mice used for photometry experiments, mice were deeply anesthetized then transcardially perfused with PBS followed by 4% PFA. Brains were removed and immersion fixed in 4% PFA overnight at 4C. Brains were transferred to 30% sucrose and allowed to equilibrate, then they were frozen and sectioned at 40 um on a cryostat (Leica). Sections were washed in 0.3% PBS-T, then incubated in a blocking solution composed of 10% normal goat serum, 0.3% PBS-T, and 3M glycine for 1 hour at room temperature. Following blocking, slices were incubated in primary antibody (Chicken a GFP, 1:1000, Abcam, ab13970) diluted in blocking buffer overnight at 4C. The next day slices were washed 3x in 0.1% PBS-T then incubated in secondary antibody (Goat a Chicken:488, 1:1000, Invitrogen, A32931) for 2 hours at room temperature. Slices were then washed 3x in 0.3% PBS-T, transferred to 0.5x PBS, and mounted on glass slides with DAPI-containing mounting media (Southern Biotech). For fluorescence imaging of DREADD-mCherry and iChloC-dsRed expression, the endogenous mCherry and dsRed fluorescence was sufficient for imaging and no antibody staining was performed. Sections were imaged on a Leica DM6000B epifluorescence microscope. For characterization of DREADD expression in mice used for behavioral experiments (**Figure 3**), three equivalent sections including the HDB/MCPO/SI complex were selected from each mouse and the region of viral expression was outlined manually in FIJI and superimposed onto the matching coronal section schematic (**Figure 3J**).

**Figure 3:**
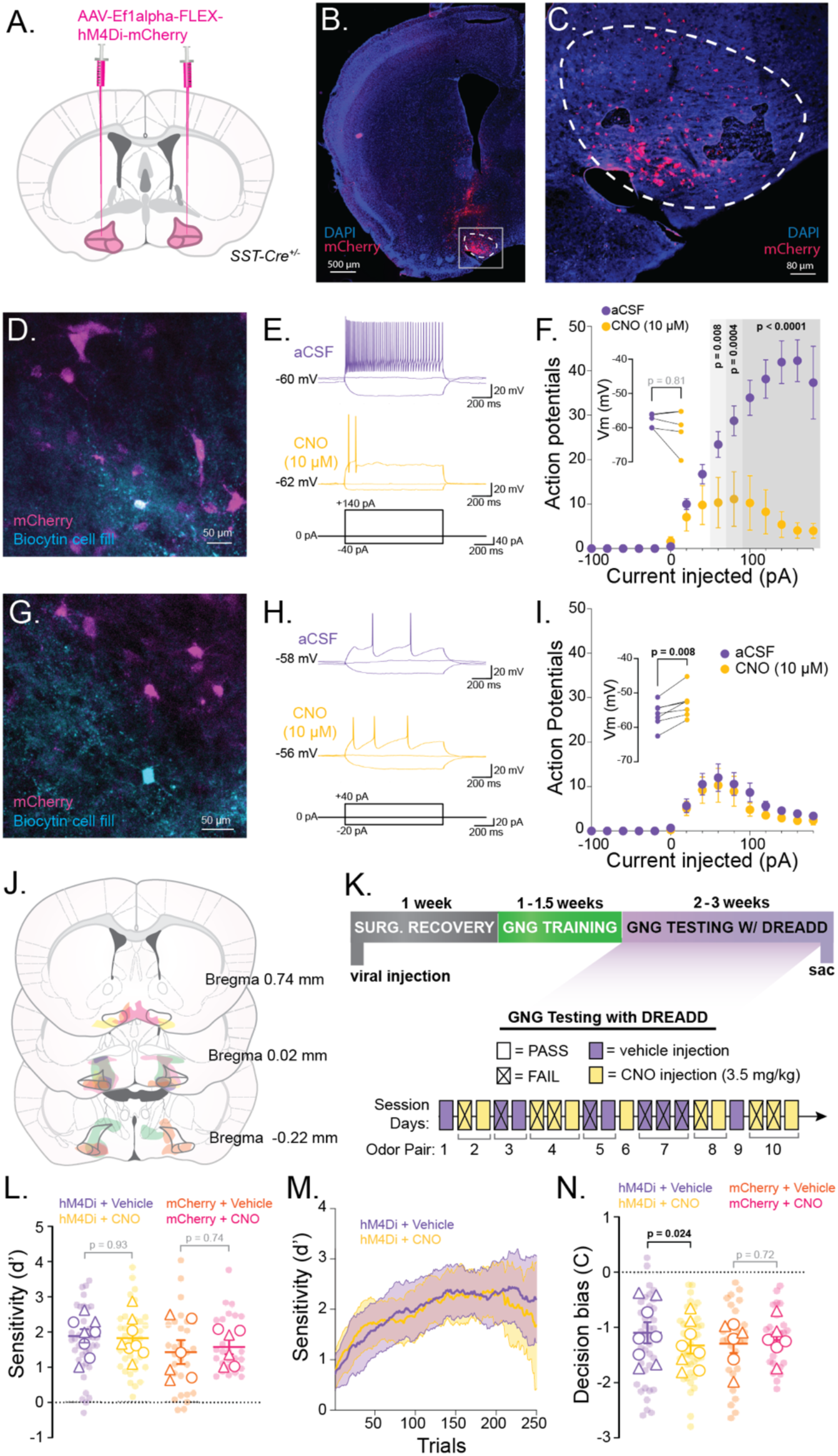
DREADD inhibition of SST neurons increases decision bias in the go/no-go task. **A.** Schematic showing bilateral injection of Cre-dependent hM4Di viral construct into basal forebrain regions including the horizontal limb of the diagonal band of Broca (HDB), magnocellular preoptic nucleus (MCPO), and substantia innominata (SI) of SST-Cre mice. **B.** Example of hM4Di-mCherry expression in BF with the HDB outlined. **C.** Inset from B showing hM4Di-mCherry expression in HDB. **D.** Example cell fill and hM4Di-mCherry expression from patched hM4Di-expressing (mCherry+) SST neuron. **E.** Example responses to current injections in aCSF (top, purple) and after CNO (10 μM) wash on (yellow middle). Bottom panel shows current injection schematic for example traces. Traces are from cell shown in D. **F.** For hM4Di-expressing neurons: Quantification of action potentials fired at each current injection in aCSF (purple) and after CNO wash on (yellow). n = 5 cells. Circles show mean ± SEM. P values from paired t tests comparing aCSF and CNO conditions at each current injection level. Inset shows resting membrane potential of each cell before (purple) and after (yellow) wash on of CNO. P value from paired t test. **G.** Example cell fill and hM4Di-mCherry expression from patched neuron not expressing hM4Di. **H.** Example responses to current injections in aCSF (top, purple) and after CNO (10 μM) wash on (yellow middle). Bottom panel shows current injection schematic for example traces. Traces are from cell shown in G. **I.** For patched neurons not expressing hM4Di: Quantification of action potentials fired at each current injection in aCSF (purple) and after CNO wash on (yellow). Circles show mean ± SEM. Inset shows resting membrane potential of each cell before (purple) and after (yellow) wash on of CNO. P value from paired t test. **J.** Quantification of DREADD expression in each animal used for behavioral experiments. Shaded regions represent areas containing cells expressing hM4Di-mCherry across three coronal sections spanning AP from bregma -0.22 – 0.74 mm. Regions are color coded by animal (N = 8). **K.** Timeline of surgery, behavioral training, and behavioral testing with DREADD-mediated inhibition (top) and schematic of the CNO or vehicle injections during testing on different odor pairs (bottom). **L.** Average d’ for individual sessions (transparent circles, n = 30-40) and mouse averages (white circles represent males and white triangles represent females) with mean and SEM of mouse averages. P values are from paired t tests between CNO and vehicle-treated sessions in DREADD-expressing experimental animals (N = 8 mice) and mCherry-expressing control animals (N = 6 mice). **M.** Performance of mice in the testing phase quantified as a rolling window of sensitivity (d’) across trials. Averages of all CNO-treatment sessions are shown in yellow with 95% confidence intervals (shaded yellow). Averages of all vehicle-treatment sessions are shown in purple with 95% confidence intervals (shaded purple). **N.** Average decision bias (C) values for individual sessions (transparent circles) and mouse averages (white circles represent males and white triangles represent females) with mean and SEM of mouse averages (n = 5 sessions per treatment per mouse). P values are from paired t tests between CNO and vehicle-treated sessions in DREADD-expressing experimental animals (N = 8 mice) and mCherry-expressing control animals (N = 6 mice).

### Experimental Design and Statistical Analyses

All statical analyses were caried out in Prism (GraphPad). Tests performed along with relevant F, t, and degrees of freedom are reported in the text. For insignificant comparisons, exact p values are reported. Significant p values are reported as p < 0.5, 0.01, 0.001, or 0.0001 as appropriate. Errors reported in the text are SEM unless otherwise noted. For comparisons across four trial types where values were matched by animal and session, repeated measures one-way ANOVAs were used with Tukey corrections for multiple comparisons. For comparisons across two trial types or groups where values were matched by animal and session, paired t tests were used. For comparisons to a null hypothesis value, one sample t tests were used. When mouse average values are reported, overall averages and errors are calculated from mouse average values. For correlation analyses, correlations are reported as the R^2^ values of linear regressions. Lines of best fit, confidence intervals, slopes, intercepts, and significance values in were calculated with simple linear regressions. For analyses comparing CNO and vehicle treatment in control and experimental animals paired t tests were performed separately for control and experimental mice to compare CNO to vehicle treated sessions. Data from odor pairs for which a mouse performed fewer than 100 total trials were excluded from all analyses. Four sessions of go/no-go behavior were excluded from analyses because of behavioral apparatus malfunctioning and three sessions were excluded for improper laser parameters.

## RESULTS

### BF PV neurons are excited during trial initiation and suppressed by reward delivery

To assess PV neuronal responses during odor discrimination learning, we selectively expressed the genetically encoded calcium indicator GCaMP8s in BF PV neurons by injecting an AAV expressing Cre-dependent GCaMP8s (AAV hsyn-FLEX-GCaMP8s) into the BF of PV-Cre mice. At the same time, we placed a fiberoptic implant over the horizontal limb of the diagonal band of Broca (HDB) in the BF (**Figure 1A**). Implant and viral targeting were verified via post hoc immunofluorescence imaging (**Figure 1B, C**. Implanted mice were then trained and tested on a go/no-go paradigm as previously described (Hanson et al., 2021). During go/no-go training, mice learned to distinguish an S+ odorant associated with a water reward from an S-odorant not associated with reward. In this odor-cued go/no-go discrimination task, there were four trial outcomes depending on the odor presented and the decision to go or not go for a reward. Retrieving the water reward upon S+ cue presentation was considered a “Hit”. Not retrieving water after S+ presentation was considered a “Miss”. Attempting to retrieve water upon S-cue presentation was a “False Alarm”. Finally, initiating a new trial after S-presentation was considered a “Correct Reject” (**Figure 1D**). Punishment for False Alarm trials consisted of a 4 second time-out period in which they were unable to initiate another trial. Mice were trained on the go/no-go task for 10-14 days prior to testing and fiber photometry recording. After training, mice were tested on novel odor pairings (**Table 1** in methods) while neuronal activity was recorded with fiber photometry. To quantify behavioral performance and decision making we used the signal detection theory measures of d’ and criterion (C). d’ measures the sensitivity of odor discrimination with higher d’ values indicating more accurate discrimination between the S+ and S-odors and chance equaling zero. Criterion measures the bias in decision-making toward reward seeking (go) or non-reward-seeking (no-go) behavior with negative C values indicating a bias towards reward-seeking behavior. Notably, in our go/no-go task, S+ and S-odors are presented with equal frequency so that optimal decision bias in the task is 0. However, mice performed this task with a significantly negative decision bias (i.e., a bias towards reward-seeking, C: -0.11 ± 0.05, p < 0.05 one sample t test) likely because the positive possibility of a water reward outweighed the negative possibility of a timeout punishment. Notably, all mice effectively learned to discriminate new pairs of S+ and S-odors with performance significantly above chance within 400 trials (d’: 2.30 ± 0.17, p < 0.0001, t = 14.56, df = 26, one-sample t test). In a separate cohort of mice, as a control for odor detection vs. operant trial initiation, we broadly expressed GCaMP in BF GABAergic neurons by injecting Cre-dependent GCaMP8s in VGAT-Cre mice. We then subjected them to a separate “go/go” behavioral task. The initial training stages for the go/go task were similar to the go/no-go task, but the S-odor was never introduced. For the go/go task, the S+ training odor was presented in half the trials, and no odor was presented in the other trials. Rewards were available after every trial regardless of whether the odor was presented.

To determine HDB PV neuronal activity across phases of the go/no-go task and different trial outcomes, photometry traces were first aligned to odor port entry times and sorted by trial type. We observed a distinct pattern of PV neuronal activity over the course of a go/no-go trial, with excitation triggered by trial initiation across all trial types and suppression following reward delivery in Hit trials (**Figure 1E**). The peak in z scored PV neuronal activity following trial initiation was equivalent across all four trial types (**Figure 1F**, Hit: 1.73 ± 0.21, Miss: 1.71 ± 0.20, Correct Reject: 1.64 ± 0.20, False Alarm: 1.68 ± 0.21 z score, p = 0.40, F = 0.93, df = 23, RM one-way ANOVA) suggesting that the response was not odor specific (S+ vs. S-), predictive of behavioral choice (go vs. no-go), or predictive of accuracy (correct vs incorrect decisions) on individual trials. Additionally, GABAergic neuronal responses to trial initiation were similar across odor and no-odor trials in the go/go behavioral task (**Figure 1F**, VGAT-Cre Go/Go Odor: 1.08 ± 0.28 z score; VGAT-Cre Go/Go No Odor: 1.13 ± 0.17 z score; p 0.84 = t = 0.21, df = 7, paired t test). These data suggest that the BF GABAergic neurons are broadly excited during operant trial initiation.

To further examine the response of PV neurons to reward seeking and reinforcement, we realigned Hit and False Alarm trials to the timing of reward port entry (**Figure 1G**). Quantifying the area under the curve (AUC) for the first 5 seconds after reward port entry showed that PV neuronal activity was strongly suppressed in Hit, but not False Alarm trials (**Figure 1H**, Hit: -2.52 ± 0.50, False Alarm: 2.60 ± 0.40 z score*s, p < 0.001, t = 6.88, df = 5, paired t test), implying that the suppression was a consequence of obtaining the water reward. Notably, this pattern of odor-evoked excitation and reward-evoked suppression mirrored the changes in both the cholinergic tone and general GABAergic neuronal activity we have previously described in the HDB (Hanson et al., 2021). These data indicate that PV neurons in the HDB respond bidirectionally during trial initiation and positive reinforcement in a go/no-go task, thus raising the question of whether PV neuronal activity controls, or is modulated by, task performance across sessions and/or learning within sessions.

To determine the extent to which PV neuronal activity was related to task performance, we examined correlations between the initial excitation during trial initiation, the reward response magnitude, behavioral performance, and decision making. Using linear regression to examine the relationship between PV neuronal activity and sensitivity across sessions, we found that d’ was negatively correlated with the trial initiation response in all trial types (**Figure 1I**, Hit slope = -0.34 ± 0.11, r^2^ = 0.16, p = 0.0046; Miss slope = -0.33 ± 0.09, r^2^ = 0.20, p = 0.0011; Correct Reject slope = -0.30 ± 0.11, r^2^ = 0.14, p = 0.0084; False Alarm slope = -0.25 ± 0.09, r^2^ = 0.13, p < 0.0107). The slopes and intercepts between trial types were not significantly different (slopes: p = 0.94, F = 0.14, df = 192, pooled slope = -0.30; intercepts: p = 0.54, F = 0.72, df = 195, pooled intercept = 2.75). These data show that in sessions where the average trial initiation response among HDB PV neurons was higher, overall performance on the discrimination task was worse. Using the same analysis to compare decision bias to PV neuronal activity, we found that C was not correlated with the magnitude of the trial initiation response for any trial type (Hit slope = -0.76 ± 0.5, r2 = 0.08, p = 0.15; Miss slope = -0.81 ± 0.4, r2 = 0.13, p = 0.07; Correct Reject slope = -0.76 ± 0.5, r2 = 0.08, p = 0.15; False Alarm slope = - 0.62 ± 0.5, r2 = 0.05, p = 0.25). The slopes and intercepts were also not different between trial types (slopes: p = .99, F = 0.03, df = 192, pooled slope = -0.74; intercepts: p = 0.94, F = 0.12, df = 195, pooled intercept = 1.87). Together these data help distinguish between neuronal activity patterns related to decision making behavior and task difficulty (i.e. the discriminability of individual odor pairs). While these data do not indicate whether the magnitude of the PV trial initiation response influences odor discrimination, they do suggest that the observed peak in HDB PV neuronal activity is related to odor discriminability, but not decision-making behavior in this go/no-go task.

Next, we examined whether PV neuronal activity in response to reward delivery was related to odor discriminability or decision-making behavior. Using the same linear regression analyses, we found no correlation between d’ and the magnitude of the reward-evoked suppression in Hit or False Alarm trials (**Figure 1J**, Hit slope = 0.23 ± 0.52, r2 = 0.004, p = 0.66; False Alarm slope = 0.64 ± 0.43, r2 = - 0.044, p = 0.14). However, we noted a significant difference between the intercepts of the regressions for Hit and False Alarm trials (Hit intercept = -2.58 ± 1.08, False Alarm intercept = 1.12 ± 0.90; p < 0.0001, F = 116.4, df = 97), confirming that the difference in AUC between Hit and False Alarm trials is consistent across sessions with a range of d’. At the same time, we found no correlation between C and the magnitude of reward evoked suppression in Hit or False Alarm trials (**Figure 1K**, Hit slope = 0.18 ± 1.24, r^2^ = 0.0004, p = 0.88; False Alarm slope = 1.34 ± 1.03, r^2^ = 0.034, p = 0.20). Intercepts were also significantly different between Hit and False Alarm trials (Hit intercept = -2.11 ± 0.33, False Alarm intercept = 2.47 ± 0.28; p < 0.0001, F = 115.5, df = 97), confirming that the reward-evoked suppression of PV neuronal activity was present across sessions with a range of decision biases.

While session averages of d’ and C help establish relationships between behavior and PV neuronal activity across sessions, we also sought to determine whether and how PV activity patterns were modulated by changes in performance over the course of a session (as performance changes with learning). To examine this, we first subdivided sessions into blocks, including the first 20 trials of an odor pair, and best blocks, including the block of 20 trials with the highest accuracy in a session. As expected, accuracy in the best blocks was consistently higher than accuracy in the first blocks, showing that mice learned to discriminate novel odor pairs over the course of the sessions (**Figure 1L**, best block – first block = 0.31 ± 0.02, p < 0.0001, t = 18.99, df = 26). Interestingly, we found that odor evoked response peaks were not different between first and best blocks (**Figure 1M**). However, the reward-evoked suppression of PV neuronal activity was greater in the best blocks compared to the first blocks (**Figure 1N, O**, best block – first block = -2.57 ± 0.37, p = 0.0010, t = 6<u>.87</u>, df = 5, paired t test). Together these findings indicate that the reward-related suppression of PV neurons, but not odor response magnitudes, changed with learning.

### SST neurons are excited during trial initiation and reward seeking

Within the BF, PV neurons receive inhibitory input from neighboring SST neurons. This disinhibitory circuit contributes to the opposing effects that BF SST and PV neurons have on arousal state (Xu et al., 2015). Given the nature of this connectivity, we next asked whether inhibition from SST neurons may contribute to the reward-evoked suppression that we observed in PV neurons. To evaluate how SST neurons selectively respond during odor discrimination learning, we again used fiber photometry during the go/no-go task. For this, we selectively expressed GCaMP in BF SST neurons and implanted a fiberoptic over the HDB **(Figure 2A, B)**. As with PV-Cre, mice reliably learned S+/S- odor-reward associations (d’ = 1.75 ± 0.26, p < 0.0001, t = 6.65, df = 9, one-sample t test). Aligning the photometry traces to odor port entry times revealed that SST neurons were excited during trial initiation (**Figure 2C**), and that this excitation was equivalent across trial types (**Figure 2D**, Hit = 1.19 ± 0.24, Miss = 1.25 ± 0.30, Correct Reject = 1.14 ± 0.22, False Alarm = 1.06 ± 0.24, p = 0.85, F = 0.39, df = 27, RM one-way ANOVA).

Responses were similar to those observed in PV neurons upon trial initiation. Notably, however, when traces were aligned to reward port entry (Figure 2E), we found that SST neurons were excited during reward-seeking and were excited more strongly when rewards were obtained in Hit trials. (**Figure 2F**, Hit AUC = 0.22 ± 0.12, False Alarm AUC = -0.17 ± 0.07, p = 0.0002, t = 7.062, df = 7, paired t test), in contrast to the suppression we observed in PV neurons. These data show that SST and PV neurons in the HDB exhibit opposing responses to reward delivery. Together, they support the hypothesis that SST and PV neurons serve opposing purposes in go/no-go task reinforcement. This raises the question of how SST neuronal activity is correlated with behavioral performance and decision-making, and to what extent the correlations oppose those observed in PV neurons.

With the goal of directly comparing the two primary GABAergic neuronal subtypes in the HDB, we examined correlations between the SST odor response magnitude, reward response magnitude, behavioral performance, and decision making. In contrast to PV neurons, we found slight positive correlations between d’ and the magnitude of SST excitation during trial initiation (Hit slope = 0.31 ± 0.08, r^2^ = 0.19, p = 0.0002, F = 15.56, df = 67; Miss slope = 0.33 ± 0.10, r^2^ = 0.14, p = 0.0015, F = 10.99, df = 67; Correct Reject slope = 0.24 ± 0.08, r^2^ = 0.12, p = 0.0037, F = 9.025, df = 67; False Alarm slope = 0.26 ± 0.09, r^2^ = 0.12, p = 0.0041), though there was no difference between trial types (slopes: p = .87, F = 0.24, df = 266, pooled slope = 0.28; intercepts: p = 0.17, F = 1.68, df = 269, pooled intercept = 1.04). We found stronger positive correlations between SST excitation during trial initiation and C for all trial types (**Figure 2G**, Hit slope = 0.80 ± 0.17, r^2^ = 0.26, p < 0.0001, F = 21.24, df = 59; Miss slope = 0.78 ± 0.21, r^2^ = 0.19, p = 0.0005, F = 13.38, df = 59; Correct Reject slope = 0.52 ± 0.18, r^2^ = 0.12, p = 0.0057, F = 8.23, df = 59; False Alarm slope = 0.65 ± 0.19, r^2^ = 0.17, p = 0.0010, F = 11.96, df = 59). The slopes and intercepts were not different between trial types (slopes: p = 0.71, F = 0.458, df = 235, pooled slope = 0.69; intercepts: p = 0.19, F = 1.62, df = 238, pooled intercept = 1.69). These data imply that a larger response of SST neurons upon trial initiation may be correlated with a more optimal decision bias (i.e. closer to zero). However, without a substantial number of sessions with decision bias above zero, we cannot determine whether the relationship would continue increasing linearly as criterion values become suboptimal in the positive direction. Therefore, it remains a question whether greater BF SST neuronal activity is related to optimal decision-making behavior or a decision bias away from reward seeking behavior.

When traces were aligned to reward port entry, we found a slight negative correlation between the magnitude of the reward-related response (AUC) and d’ across sessions in Hit but not False Alarm trials (Hit slope = -0.20 ± 0.09, r ^2^= 0.14, p = 0.027, F = 5.36, df = 34; False Alarm slope = 0.005 ± 0.09, r^2^ = 0.14, p = 0.95, F = 0.004, df = 34). Taken together with our finding that reward-related excitation of SST neurons is larger in Hit trials than in False Alarm trials, this analysis suggested that the difference between Hit and False Alarm trials was greatest in sessions with lower overall d’ values (indicating worse task performance). Comparing reward response magnitudes to decision bias, we found no correlation between C and the magnitude of the reward response for Hit or False Alarm trials (Hit slope = -0.15 ± 0.12, r^2^ = 0.01, p 0.42, F = 0.66, df = 56; False Alarm slope = 0.22 ± 0.16, r^2^ = 0.03, p = 0.17, F = 1.96, df = 56). However, we did observe a significant difference between the elevations of the regressions for Hit vs. False Alarm trials (**Figure 2H**, p 0.012, F = 6.53, df = 113), confirming that reward responses are greater in Hit trials compared to False Alarm trials.

Next, to determine how patterns of SST neuronal activity were modulated during the learning of a new S+/S- odor pair, we examined how odor and reward responses changed within go/no-go sessions, as mice improved at the discrimination task. To examine performance-related changes, we subdivided sessions into first blocks and best blocks (as described for PV neuronal recordings), with first blocks showing significantly lower accuracy (**Figure 2I**, Best block – first block = 0.31 ± 0.02, p < 0.0001, t = 14.8, df = 30, paired t test). Aligning first and best block Hit trials to trial initiation revealed no differences in odor-evoked peaks in SST neuronal activity (**Figure 2J**), suggesting that SST odor responses are determined by the identity of the odor pair, and are not modulated by learning. However, aligning Hit trials to reward port entry revealed a reduction in reward-related excitation of SST neurons in the best blocks compared to the first blocks (**Figure 2K, l**, best block – first block = -2.83 ± 1.15, p = 0.0434, t = 2.46, df = 7, paired t test). Taken together with the negative correlation between session average d’ values and reward-related excitation of SST neurons, these data suggest SST reward responses are modulated by performance, with improvements in discrimination leading to a reduction in reward-related excitation both across odor pairs with a range of difficulties, and within sessions, as mice learn to discriminate new S+ and S- odor pairs.

### DREADD-mediated inhibition of SST neurons does not impact firing of neighboring BF neurons

We next sought to determine how directly manipulating BF GABAergic signaling impacted odor discrimination and decision-making behavior. Towards this, we selectively expressed the inhibitory DREADD hM4Di in BF SST neurons, first confirming that treatment with the DREADD ligand CNO showed targeted and ligand-dependent inhibition (**Figure 3A-C**). We chose SST neurons for manipulation because they have been shown previously to make local inhibitory synapses onto each of the other major cell types within the HDB, including PV GABAergic, glutamatergic, and cholinergic neurons (Zaborszky and Duque, 2000; Xu et al., 2015). To determine the direct effect of CNO treatment on hM4Di expressing SST neurons, we made acute brain slices harboring the HDB for whole cell electrophysiological recordings. After current clamping mCherry labeled, hM4Di-expressing neurons (**Figure 3D**) we made stepped current injections from -100 to 180 pA before and after CNO application (10 μM). Prior to CNO, SST neurons showed fast-spiking patterns with narrow action potentials as previously described (Anaclet et al., 2018) (**Figure 3E**). As expected, we found that CNO significantly inhibited action potential firing in hM4Di expressing HDB SST neurons at current steps of 60 pA or greater (**Figure 3F**), though we did not observe systematic hyperpolarization with CNO (**Figure 3F, inset,** ΔVm -2.18 ± 1.95, p = 0.33, t = -1.12, df = 4, paired t test). To examine whether inhibition of SST neurons impacted firing properties of neighboring HDB neurons, we also patched and current clamped unlabeled, non-hM4Di-expressing neurons (**Figure 3G**). Unlabeled neurons were slow spiking and susceptible to depolarization block, suggesting that they represented cholinergic or glutamatergic subtypes (**Figure 3H**). We found that CNO treatment neither reduced, nor enhanced the firing of the non-hM4Di expressing neurons in the HDB (**Figure 3I**). CNO application did, however, cause depolarization in non-hM4Di-expressing neurons (**Figure 3I, inset,** ΔVm 3.22 ± 0.73, p = 0.0045, t = 4.41, df = 6, paired t test). Notably, the observed depolarization did not alter the overall firing properties of non-hM4Di-expressing neurons. While these data indicate that reducing the ability of SST neurons to fire action potentials does not significantly impact the ability of the other HDB neurons to fire action potentials when driven, they do not show how DREADD-mediated inhibition of SST neurons impacts HDB circuit activity *in vivo* or during behavior.

### DREADD inhibition of SST neurons modestly increases decision bias in go/no-go task

To directly examine the role of BF SST neurons in odor discrimination and decision-making behavior, we next selectively targeted hM4Di expression to BF SST neurons in the HDB, the magnocellular preoptic nucleus (MCPO), and substantia innominata (SI) – regions forming a complex within the BF that is densely interconnected with olfactory brain regions (Zheng et al., 2018) (**Figure 3J**). We then trained mice on the go/no-go odor discrimination task and tested how they performed in sessions with or without CNO treatment. In a subset of mice, mCherry instead of hM4Di-mCherry was selectively expressed in SST neurons to serve as controls for CNO treatment. After training, mice were tested on new S+/S- odor pairs for consecutive days until they either (1) successfully learned the new odor associations (achieved two consecutive blocks of 85% accuracy), or (2) failed to meet the criteria for learning three days in a row. Odor pairs were then chosen to span a range of difficulties and were approximately matched in difficulty between CNO and Vehicle treated sets of odors (Table 2 in methods). Mice were treated with either CNO (3.5 mg/kg) or Vehicle (1% DMSO in saline) for each day of alternating odor pairs (**Figure 3K**). Sessions across multiple days of the same odor pair received the same CNO or Vehicle treatment and were concatenated into single sessions from which sensitivity and decision bias measures (d’ and C) were calculated. Despite our earlier finding that discrimination sensitivity was inversely correlated with reward related excitation in SST neurons, we found that inhibiting SST neurons did not impact odor discrimination at the level of whole sessions (**Figure 3L**, Hm4Di CNO-Vehicle = -0.06 ± 0.0.18, p = 0.74, t = 0.34, df = 7; Control CNO-Vehicle = 0.14 ± 0.0.19, p = 0.49, t = 0.75, df = 5, paired t tests). Additionally, inhibiting SST neurons did not impact the rate at which mice learned new odor associations (**Figure 3M**). These data suggest that odor and reward evoked excitation of HDB SST neurons does not directly impact odor discrimination or odor discrimination learning. Notably, however, we found that decision bias was subtly but significantly altered by CNO treatment in hM4Di expressing mice (**Figure 3N**, Hm4Di CNO-Vehicle = -0.11 ± 0.03, p < 0.05, t = 3.38, df = 7; Control CNO-Vehicle = 0.04 ± 0.05, p = 0.47, t = 0.78, df = 5, paired t tests). Thus, when HDB SST neurons were inhibited, mice showed a slightly stronger bias toward reward seeking behavior (i.e. a more negative C value). The bias towards reward seeking moved decision-making further away from the optimal decision bias of zero for our go/no-go task. Together, these data suggest that HDB SST may affect reward-seeking without directly impacting odor discrimination or learning in the context of a freely moving, self-initiated, go/no-go discrimination task.

### Optogenetic activation of PV neurons after reward delivery does not influence odor discrimination or decision bias

To investigate the role of BF PV neurons in odor discrimination and decision-making, we targeted conditional ChR2-GFP (AAV ef1a-Flex-hChR2-eYFP) to the BF neurons in the HDB of PV-cre mice and placed fiber optic implants above the region to selectively activate the PV neurons (**Figure 4A-C**). Mice were then trained on the go/no-go odor discrimination task while tethered to an optogenetic cable. After training, mice received one of three optogenetic stimulation conditions during different go/no-go testing sessions with novel S+/S- odor pairs (**Figure 4D**, **Table 3**). In the first condition, mice received 5 seconds of 20 Hz stimulation each time they received a reward on a Hit trial (**Figure 4D**, reward-locked stimulation). In the second condition, mice received 5 seconds of 20 Hz stimulation every 20 seconds regardless of where they were in a behavioral trial (**Figure 4D**, regular stimulation). In the last control condition, mice were tethered to a fiberoptic cable but received no stimulation (**Figure 4D**, no stimulation). Mice were randomized to different combinations of stimulus conditions and odor pairs. Optogenetic stimulation time locked to reward delivery, during a time when PV neuronal activity is suppressed **(Figure 1 G, H)**, did not impact the overall accuracy of odor discrimination compared to (**Figure 4E**, p = 0.19, F = 1.97, df = 26, RM one-way ANOVA). Optogenetic stimulation also did not affect decision-bias, as C values across the three stimulation types showed no significant difference (**Figure 4F**, p = 0.14, F = 2.23, df = 26, RM one-way ANOVA).

**Figure 4:**
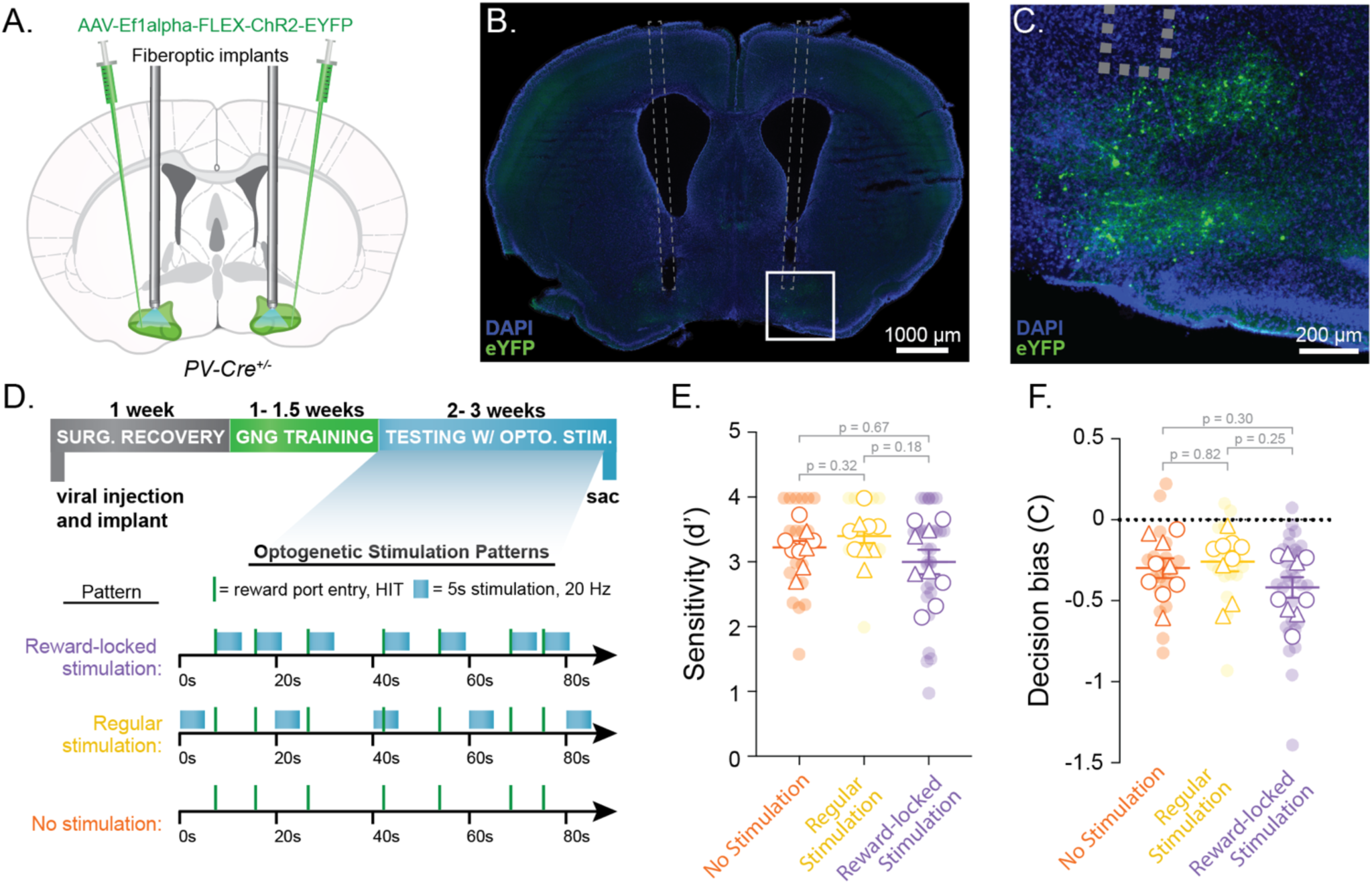
Optogenetic stimulation of PV neurons during reward delivery does not affect go/no- go task performance. **A.** Schematic showing bilateral injection of Cre-dependent ChR2 viral construct and fiberoptic implant in PV-Cre mice. Green-highlighted regions of the BF include the horizontal limb of the diagonal band of Broca (HDB), the magnocellular preoptic nucleus (MCPO) and substantia innominata (SI). **B.** Example slice image showing the implant tract, expression of ChR2-eYFP (green), and DAPI (blue). **C.** Viral expression (green) and implant tip localization (dashed line) in the HDB. **D.** Timeline of surgery, behavioral training, and behavioral testing with optogenetic stimulation (top) and schematic outlining the different optogenetic stimulation experimental conditions (bottom). **E.** Summary d’ for individual sessions, calculated as the maximum d’ from a 40-trial rolling window over the course of a session (see methods) (transparent circles, n = 19 no stimulation sessions, 18 regular stimulation sessions, and 30 reward-locked stimulation sessions). Mouse averages of summary d’ values are shown as white circles for males and white triangles for females with mean and SEM (N = 9 mice, p values from RM one-way ANOVA with multiple comparisons using Tukey correction). **F.** Average criterion values for individual sessions (transparent circles) and mouse averages (white circles for male and white triangles for females) with mean and SEM of mouse averages. (p values from RM one-way ANOVA with multiple comparisons using Tukey correction).

### Optogenetic inhibition of PV neurons upon trial initiation improves odor discrimination without impacting decision bias

To determine the impact of BF PV excitation upon trial initiation on odor discrimination, we next targeted the inhibitory light activated chloride channel iChloC (AAV-ef1a-Flex-iChloC-2A-dsRed) to BF PV neurons (**Figure 5**). To confirm the extent and time course of iChloC-mediated inhibition, we made acute brain slices containing HDB and performed loose cell attached electrophysiological recordings from dsRed+ BF PV neurons (**Figure 5A**). In dsRed + cells, a 500 ms pulse of blue light produced a substantial reduction in firing relative to baseline which persisted for several seconds after the offset of the light pulse **(Figure 5B, C)**. By 10 seconds after light offset, baseline firing had recovered fully (**Figure 5D**, magenta circles, ΔPre vs. Light on normalized firing rate = -0.93 ± 0.04, p < 0.0001, q = 29.94, df = 4; ΔPre vs. 10 s Post normalized firing rate = -0.01 ± 0.09, p = 0.9914, q = 0.18, df = 4; ΔLight on vs. Post normalized firing rate = 0.92 ± 0.10, p = 0.0015, q = 13.41, df = 4, RM one-way ANOVA). There was no impact of light on neighboring dsRed – neurons (**Figure 5D**, grey circles, ΔPre vs. Light on normalized firing rate = 0.05 ± 0.04, p 0.50, q = 1.90, df = 2; ΔPre vs. 10 s Post normalized firing rate = 0.01 ± 0.06, p = 0.9936, q = 0.15, df = 2; ΔLight on vs. Post normalized firing rate = -0.04 ± 0.07, p = 0.84, q = 0.83, df = 2, RM one-way ANOVA).

**Figure 5:**
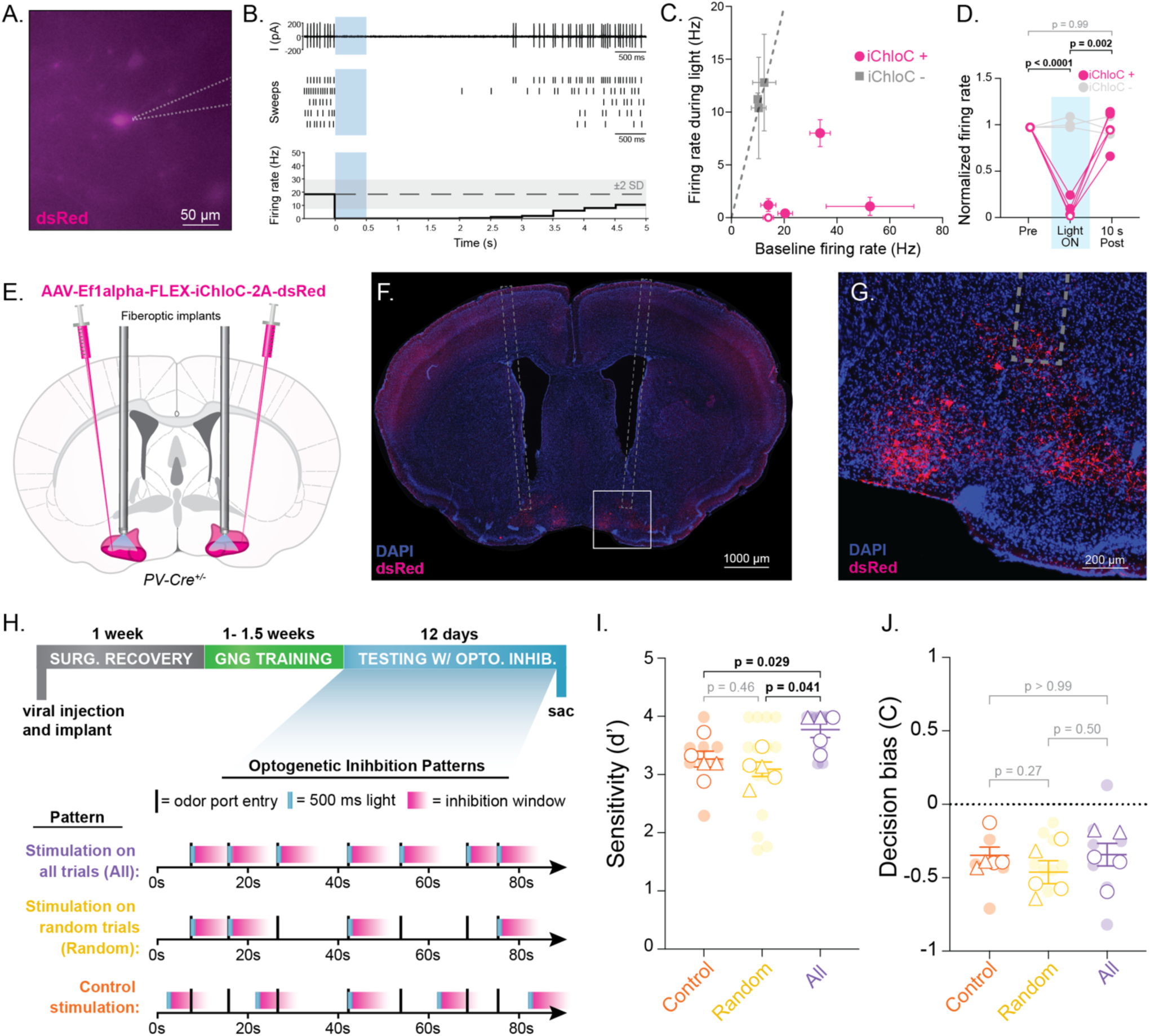
Optogenetic inhibition of PV neurons during odor presentations does not affect go/no-go task performance. **A.** ds-Red fluorescence of iChloC-expressing PV neuron during loose cell attached electrophysiological recording in slice. Grey dashes show electrode outline. **B.** Example iChloC+ cell firing in loose cell attached configuration before, during, and after a 500 ms pulse of LED light (transparent blue block). Top panel shows an example trace. Middle panel shows raster plot of five trials. Bottom panel shows the normalized average firing rate across sweeps, binned by 500 ms. The range within two standard deviations of the baseline firing rate is shown as shaded grey. **C.** Baseline firing rate versus firing rate during light pulses for each iChloC+ (magenta) and iChloC-cell (grey). Error bars are SEM of baseline firing rate (along X axis) and SEM of firing rate during the light pulse (along Y axis). Identity line is shown as grey dashes. White marker indicates example neuron shown in panels A and B. **D.** Quantification of normalized firing rates before light, during light, and 10 seconds after light for iChloC+ (magenta) and iChloC-cells (grey). P values are from RM one-way ANOVA of iChloC+ neurons corrected for multiple comparisons. P values comparing iChloC-cells are insignificant and not shown. White markers indicate example neuron shown in panels A and B. **E.** Schematic showing bilateral injection of Cre-dependent iChloC viral construct and fiberoptic implant in PV-Cre mice. Red-highlighted regions of the BF include the horizontal limb of the diagonal band of Broca (HDB), the magnocellular preoptic nucleus (MCPO) and substantia innominata (SI). **F.** Example slice image showing the implant tract and expression of iChloC-dsRed. **G.** Viral expression and implant tip localization in the HDB. **H.** Timeline of surgery, behavioral training, and behavioral testing with optogenetic inhibition (top) and schematic outlining the different optogenetic inhibition experimental conditions (bottom). **I.** Summary d’ for individual sessions, calculated as the maximum d’ from a 40-trial rolling window over the course of a session (transparent circles, n = 8 control sessions, 14 random trial stimulation sessions, and 10 all trial stimulation sessions). Mouse averages of summary d’ values are shown as white circles (males) and white triangles (females) with mean and SEM. P values are from RM one-way ANOVA with multiple comparisons using Tukey correction. **J.** Average criterion values for individual sessions (transparent circles) and mouse averages (white circles and triangles) with mean and SEM of mouse averages. P values are from RM one-way ANOVA with multiple comparisons using Tukey correction.

For *in vivo* inhibition of BF PV neurons during behavior, we placed bilateral fiberoptics over the HDB (**Figure 5E-G**) and trained mice to perform the go/no-go task. After training, mice received one of three optogenetic stimulation conditions (**Figure 5H**) during different go/no-go testing sessions with the same set of S+/S- odor pairs used in stimulation experiments (**Table 3**). In the first condition, mice received 500 ms of continuous stimulation upon entering the odor port on every trial (**Figure 5H**, All). In the second condition, mice received 500 ms of continuous stimulation on 50% of the trials selected at random (**Figure 5H**, Random). In the third condition, mice received 500 ms of continuous stimulation every 20 seconds regardless, analogous to the “regular stimulation” condition for optogenetic stimulation (**Figure 5H**, Control). Mice were randomized to different combinations of stimulus conditions and odor pairs. Surprisingly, we found that optogenetically inhibiting BF PV neurons during all odor presentations within a session improved odor discrimination (**Figure 5I**). D’ values were significantly higher in sessions where BF PV neurons were inhibited during all odor presentations compared to sessions where only 50% of trials triggered inhibition and sessions where inhibition was not time-locked to odor presentations (p = 0.0089, F = 11.8, df = 14, RM one-way ANOVA). Inhibition of BF PV neurons did not, however, affect decision-bias (**Figure 5J**, p = 0.35, F = 1.18, df = 14, RM one-way ANOVA) indicating that the reduction in errors was balanced across miss and false alarm-type errors. In the random condition, within sessions, mice performed with similar d’ and C on inhibited and uninhibited trials (d’: p = 0.40, t = 0.87, df = 13; C: p = 0.68, t = 0.42, df = 13, paired t tests). This suggests that the impact of inhibition on accuracy was due to a cumulative effect across the session rather than an increase in discriminatory ability on individual trials. Together, these results highlight distinct influences of BF PV neurons on odor discrimination learning and BF SST neurons on odor-guided decision-making behavior.

## DISCUSSION

Basal forebrain (BF) cholinergic neurons play a critical role in regulating behavioral states, mediating attention and arousal, and are vulnerable to neurodegenerative diseases like Alzheimer’s and Parkinson’s (Sarter and Bruno, 1997; Levin and Simon, 1998; Sarter et al., 2001; Grothe et al., 2010; Parrao et al., 2012). These neurons respond to sensory cues, reinforcement, and shifts in behavioral state across various timescales (Phillis, 1968; Szymusiak, 1995; Sarter and Bruno, 1999; Hasselmo and McGaughy, 2004; Parikh et al., 2007; Chaudhury et al., 2009; Hangya et al., 2015; Xu et al., 2015; Ruivo et al., 2017; Patel et al., 2019). Similarly, non-cholinergic BF neurons are strongly associated with sensory perception and behavioral task performance, exhibiting activity changes in response to conditioned stimuli and reinforcement (Lin and Nicolelis, 2008; Hassani et al., 2009; Devore et al., 2015; Hangya et al., 2015; Harrison et al., 2016; Zheng et al., 2018; Nunez-Parra et al., 2020). BF neuronal activity is also modulated by expectation, with more surprising reinforcement eliciting larger increases in firing for both cholinergic and non-cholinergic BF neurons (Lin and Nicolelis, 2008; Hangya et al., 2015; Zheng et al., 2018).

In the olfactory system, BF cholinergic and GABAergic neurons project extensively to the olfactory bulb (OB) (Zaborszky et al., 1986). GABAergic projections preferentially target specific GABAergic interneurons in the OB, where they influence fine odor discrimination, signal to noise ratios, and the survival of adult born neurons (Böhm et al., 2020; Hanson et al., 2020; De Saint Jan, 2022). A direct comparison of the impact of BF cholinergic and GABAergic projections on OB activity reveals that BF GABAergic neurons preferentially increase odor-evoked mitral cell firing and suppress spontaneous activity, while cholinergic neurons increase spontaneous, sniff-locked, and odor evoked mitral cell firing (Böhm et al., 2020). In olfactory tasks, cholinergic and non-cholinergic BF neurons respond to conditioned stimuli, unconditioned stimuli, and both positive and negative reinforcement (Hangya et al., 2015; Nunez-Parra et al., 2020; Hanson et al., 2021), with a significant proportion of BF neurons showing suppression in response to odor stimuli and positive reinforcement (Nunez-Parra et al., 2020). Additionally, we have recently shown that BF GABAergic activity is powerfully suppressed after positive reinforcement (Hanson et al., 2021). Previous studies, however, did not differentiate between subtypes of non-cholinergic neurons and were, therefore, unable to discern distinct roles of BF GABAergic neuronal subtypes on odor-guided behavior.

Here, we targeted PV and SST GABAergic neurons in the BF to uncover their distinct contributions to olfactory discrimination and decision-making. Both subtypes were excited during operant trial initiation, but their responses to positive reinforcement diverged – SST neurons showed transient excitation, while PV neurons exhibited sustained suppression. These opposing responses may reflect direct inhibition of PV neurons by SST neurons, though the congruency of their activity during trial initiation suggest that this relationship is more complicated. It is possible, for example, that the inhibitory input from SST to PV neurons is gated differently in response to external sensory inputs than it is for internal state and/or reinforcement information. While our findings do not suggest specific gating mechanisms, they do reveal a clear divergence in how BF neurons are recruited during operant trial initiation and in response to reinforcement feedback.

The opposing reward-related responses of PV and SST neurons parallel their roles in behavioral state regulation, suggesting they influence discrimination learning and decision-making differently (Xu et al., 2015; Harrison et al., 2016). Correlational analyses provide insights into their functions but cannot determine causality, as observed activity could result from neural control of behavior, behavioral feedback, or overarching state changes. Along these lines, our finding that GABAergic neurons, targeted with VGAT-Cre, are excited during trial initiation regardless of whether an odor is presented suggests that BF PV and SST neuronal activity is driven by operant behavior – an example of behavior or behavioral state driving neural activity. At the same time, our findings that manipulating PV and SST neuronal activity alter decision making and discrimination accuracy provide examples of neural activity driving behavior. Together, across our odor-guided behavioral task, we find that BF neurons respond to external cues (e.g., reinforcement), internal state signals (e.g., trial initiation), and influence behavioral performance (e.g., bias and accuracy). Thus, our results demonstrate a complex interplay between behavior, behavioral state, odor perception, and BF GABAergic neuronal activity that varies across the stages of an odor-guided behavioral task in a cell type-specific manor.

SST neurons are intriguing components of the BF circuit because they directly inhibit each of the other major cell types in the BF, including glutamatergic, cholinergic, and PV GABAergic neurons. To help determine the impact of SST neuronal excitability on other cell types of the BF, we expressed an inhibitory DREADD in SST neurons and measured action potential firing and resting membrane potentials in both DREADD and non-DREADD expressing neurons. While CNO treatment effectively inhibited action potential firing of SST neurons driven by current injections, it did not systematically alter action potential firing in neighboring non-DREADD expressing neurons. However, CNO treatment did lead to a slight increase in the resting membrane potential of non-DREADD expressing neurons. This suggests that SST neurons may be tonically inhibiting neighboring BF neurons, and that DREADD-mediated inhibition of SST neurons relieves some of this inhibition, raising the resting potential of neighboring neurons.

To investigate whether SST excitation, either upon trial initiation or with reward-seeking was having an impact on decision making behavior, we chemogenetically inhibited SST neurons during a subset of go/no-go testing sessions. Chemogenetic inhibition had the advantage of reliably reducing both trial initiation-linked and reward-linked inhibition across all trials of a behavioral session while also reducing tonic inhibition of neighboring BF neurons, as demonstrated with slice electrophysiology. Based on our observed correlations, we predicted that (1) if SST excitation during odor exposure was driving more optimal decision-making behavior, then inhibiting SST neurons would lead to larger (more negative) decision biases, and (2) if a reduction in SST reward-related excitation facilitated learning, then inhibiting SST neurons would increase the rate at which discrimination sensitivity (d’) improved within sessions. Comparing go/no-go discrimination testing sessions in which BF SST neurons were chemogenetically inhibited to sessions in which BF SST neurons were not inhibited, we found that DREADD-mediated inhibition of SST neurons led to a modest reduction in C with no change in discrimination within or across sessions. Together, these data suggest that SST neuronal activity influences reward-seeking behavior as predicted without altering discrimination ability or discrimination learning directly.

Finally, to investigate whether PV neuronal was driving decision-making behavior or influencing odor discrimination, we incorporated optogenetic manipulations of BF PV neurons into the go/no-go assay. For these experiments, temporally specific optogenetic stimulation allowed us to restrict excitation to the period immediately following reward delivery, thus allowing us to test whether the specificity of PV suppression after reward delivery was necessary for odor discrimination learning. Although we had previously found that suppression of PV neuronal activity was correlated with reward delivery, optogenetic stimulation of the BF PV neurons to reduce the reward-linked suppression did not affect either decision-bias or odor discrimination. Instead, optogenetic inhibition of PV neurons during odor presentations (blunting BF PV excitation upon trial initiation and driving lasting suppression regardless of reward status) effectively increased accuracy without affecting decision bias. The choice of optogenetic, trial-by-trial inhibition over DREADD-mediated whole session inhibition introduced variability within behavioral sessions but allowed us to compare subsets of inhibited to uninhibited trials within individual sessions. Notably, when PV neurons were inhibited on a subset of trials, accuracy was similar across inhibited and uninhibited trials, suggesting that the impact of PV inhibition on discrimination accuracy is cumulative across trials – likely impacting discrimination learning over the course of a session rather than acute discrimination on individual trials. These data indicate that inhibiting BF PV neurons improves odor discrimination learning, and they agree with the negative correlation we observed between session d’ averages and PV excitation upon trial initiation (**Figure 1I**). Optogenetic inhibition of PV neurons with iChloC also led to lasting suppression of PV neurons into the post-reward/reward-seeking phase of the task. Therefore, our optogenetic inhibition paradigm led to artificial suppression of PV neurons after incorrect reward seeking on False Alarm trials. Our finding that discrimination accuracy, nevertheless, increased under these conditions suggests that specific, reward-linked suppression of PV neurons is not necessary for discrimination learning. Together with the correlation analyses, our optogenetic inhibition results suggest that excitation of BF PV neurons during odor detection, but not their suppression after reward delivery drives odor discrimination learning.

In conclusion, our findings reveal distinct roles for BF PV and SST neurons in an odor-guided task. Specifically, we demonstrate similar recruitment of BF SST and PV neurons during the internally driven initiation of an operant behavior, but opposite responses to external, rewarding reinforcement. Furthermore, we demonstrate distinct influences of BF SST neurons on decision-making behavior and BF PV neurons on discrimination learning. Our results advance our understanding of how specific BF GABAergic subtypes integrate sensory and behavioral information, shedding light on the neural mechanisms underlying sensory perception, decision-making, and learning.

## Acknowledgements

This work was supported by the NIH through NINDS and the BRAIN Initiative (U01NS111692 and R01NS078294 to BRA) and NIDCD and the BRAIN Initiative (K99DC019505 to EHM).

## REFERENCES

Anaclet C, De Luca R, Venner A, Malyshevskaya O, Lazarus M, Arrigoni E, Fuller PM (2018) Genetic activation, inactivation, and deletion reveal a limited and nuanced role for somatostatin-containing basal forebrain neurons in behavioral state control. Journal of Neuroscience 38:5168–5181.

Anaclet C, Pedersen NP, Ferrari LL, Venner A, Bass CE, Arrigoni E, Fuller PM (2015) Basal forebrain control of wakefulness and cortical rhythms. Nature communications 6:8744.

Ballinger EC, Ananth M, Talmage DA, Role LW (2016) Basal forebrain cholinergic circuits and signaling in cognition and cognitive decline. Neuron 91:1199–1218.

Bendahmane M, Ogg MC, Ennis M, Fletcher ML (2016) Increased olfactory bulb acetylcholine bi-directionally modulates glomerular odor sensitivity. Scientific reports 6:1–13.

Bigl V, Woolf NJ, Butcher LL (1982) Cholinergic projections from the basal forebrain to frontal, parietal, temporal, occipital, and cingulate cortices: a combined fluorescent tracer and acetylcholinesterase analysis. Brain research bulletin 8:727–749.

Böhm E, Brunert D, Rothermel M (2020) Input dependent modulation of olfactory bulb activity by HDB GABAergic projections. Scientific reports 10:1–15.

Chaudhury D, Escanilla O, Linster C (2009) Bulbar acetylcholine enhances neural and perceptual odor discrimination. Journal of Neuroscience 29:52–60.

Chaves-Coira I, Barros-Zulaica N, Rodrigo-Angulo M, Núñez Á (2016) Modulation of specific sensory cortical areas by segregated basal forebrain cholinergic neurons demonstrated by neuronal tracing and optogenetic stimulation in mice. Frontiers in neural circuits 10:28.

De Saint Jan D (2022) Target-specific control of olfactory bulb periglomerular cells by GABAergic and cholinergic basal forebrain inputs. Elife 11:e71965.

Devore S, Pender-Morris N, Dean O, Smith D, Linster C (2015) Basal forebrain dynamics during nonassociative and associative olfactory learning. Journal of neurophysiology 115:423–433.

Gielow MR, Zaborszky L (2017) The input-output relationship of the cholinergic basal forebrain. Cell reports 18:1817–1830.

Gritton HJ, Howe WM, Mallory CS, Hetrick VL, Berke JD, Sarter M (2016) Cortical cholinergic signaling controls the detection of cues. Proceedings of the National Academy of Sciences 113:E1089– E1097.

Grothe M, Zaborszky L, Atienza M, Gil-Neciga E, Rodriguez-Romero R, Teipel SJ, Amunts K, Suarez-Gonzalez A, Cantero JL (2010) Reduction of basal forebrain cholinergic system parallels cognitive impairment in patients at high risk of developing Alzheimer’s disease. Cerebral Cortex 20:1685–1695.

Hangya B, Ranade SP, Lorenc M, Kepecs A (2015) Central cholinergic neurons are rapidly recruited by reinforcement feedback. Cell 162:1155–1168.

Hanson E, Brandel-Ankrapp KL, Arenkiel BR (2021) Dynamic cholinergic tone in the basal forebrain reflects reward-seeking and reinforcement during olfactory behavior. Frontiers in Cellular Neuroscience 15:635837.

Hanson E, Swanson J, Arenkiel BR (2020) GABAergic input from the basal forebrain promotes the survival of adult-born neurons in the mouse olfactory bulb. Frontiers in Neural Circuits 14:17.

Harrison TC, Pinto L, Brock JR, Dan Y (2016) Calcium imaging of basal forebrain activity during innate and learned behaviors. Frontiers in neural circuits 10:36.

Hassani OK, Lee MG, Henny P, Jones BE (2009) Discharge profiles of identified GABAergic in comparison to cholinergic and putative glutamatergic basal forebrain neurons across the sleep– wake cycle. Journal of Neuroscience 29:11828–11840.

Hasselmo ME, McGaughy J (2004) High acetylcholine levels set circuit dynamics for attention and encoding and low acetylcholine levels set dynamics for consolidation. Progress in brain research 145:207–231.

Herman AM, Ortiz-Guzman J, Kochukov M, Herman I, Quast KB, Patel JM, Tepe B, Carlson JC, Ung K, Selever J (2016) A cholinergic basal forebrain feeding circuit modulates appetite suppression. Nature 538:253–256.

Herrero JL, Roberts MJ, Delicato LS, Gieselmann MA, Dayan P, Thiele A (2008) Acetylcholine contributes through muscarinic receptors to attentional modulation in V1. Nature 454:1110– 1114.

Lau B, Salzman CD (2008) Noncholinergic neurons in the basal forebrain: often neglected but motivationally salient. Neuron 59:6–8.

Ledbetter NM, Chen CD, Monosov IE (2016) Multiple mechanisms for processing reward uncertainty in the primate basal forebrain. Journal of Neuroscience 36:7852–7864.

Levin ED, Simon BB (1998) Nicotinic acetylcholine involvement in cognitive function in animals. Psychopharmacology 138:217–230.

Lin S-C, Nicolelis MAL (2008) Neuronal ensemble bursting in the basal forebrain encodes salience irrespective of valence. Neuron 59:138–149.

Muir JL, Page KJ, Sirinathsinghji DJS, Robbins TW, Everitt BJ (1993) Excitotoxic lesions of basal forebrain cholinergic neurons: effects on learning, memory and attention. Behavioural brain research 57:123–131.

Nunez-Parra A, Maurer RK, Krahe K, Smith RS, Araneda RC (2013) Disruption of centrifugal inhibition to olfactory bulb granule cells impairs olfactory discrimination. Proceedings of the National Academy of Sciences 110:14777–14782.

Nunez-Parra A, Rio C-D, Christian A, Huntsman MM, Restrepo D (2020) The basal forebrain modulates neuronal response in an active olfactory discrimination task. Frontiers in Cellular Neuroscience 14:141.

Ogg MC, Ross JM, Bendahmane M, Fletcher ML (2018) Olfactory bulb acetylcholine release dishabituates odor responses and reinstates odor investigation. Nature communications 9:1–11.

Parikh V, Kozak R, Martinez V, Sarter M (2007) Prefrontal acetylcholine release controls cue detection on multiple timescales. Neuron 56:141–154.

Parrao T, Chana P, Venegas P, Behrens MI, Aylwin ML (2012) Olfactory deficits and cognitive dysfunction in Parkinson’s disease. Neurodegenerative Diseases 10:179–182.

Patel JM, Swanson J, Ung K, Herman A, Hanson E, Ortiz-Guzman J, Selever J, Tong Q, Arenkiel BR (2019) Sensory perception drives food avoidance through excitatory basal forebrain circuits. eLife 8.

Phillis JW (1968) Acetylcholine release from the cerebral cortex: its role in cortical arousal. Brain research 7:378–389.

Pinto L, Goard MJ, Estandian D, Xu M, Kwan AC, Lee S-H, Harrison TC, Feng G, Dan Y (2013) Fast modulation of visual perception by basal forebrain cholinergic neurons. Nature neuroscience 16:1857–1863.

Ruivo LMT-G, Baker KL, Conway MW, Kinsley PJ, Gilmour G, Phillips KG, Isaac JTR, Lowry JP, Mellor JR (2017) Coordinated acetylcholine release in prefrontal cortex and hippocampus is associated with arousal and reward on distinct timescales. Cell reports 18:905–917.

Sanz Diez A, Najac M, De Saint Jan D (2019) Basal forebrain GABAergic innervation of olfactory bulb periglomerular interneurons. The Journal of Physiology 597:2547–2563.

Sarter M, Bruno JP (1997) Cognitive functions of cortical acetylcholine: toward a unifying hypothesis. Brain research reviews 23:28–46.

Sarter M, Bruno JP (1999) Cortical cholinergic inputs mediating arousal, attentional processing and dreaming: differential afferent regulation of the basal forebrain by telencephalic and brainstem afferents. Neuroscience 95:933–952.

Sarter M, Givens B, Bruno JP (2001) The cognitive neuroscience of sustained attention: where top-down meets bottom-up. Brain research reviews 35:146–160.

Semba K (1991) The cholinergic basal forebrain: a critical role in cortical arousal. The basal forebrain:197–218.

Szymusiak R (1995) Magnocellular nuclei of the basal forebrain: substrates of sleep and arousal regulation. Sleep 18:478–500.

Szymusiak R, Alam N, McGinty D (2000) Discharge patterns of neurons in cholinergic regions of the basal forebrain during waking and sleep. Behavioural brain research 115:171–182.

Tashakori-Sabzevar F, Ward RD (2018) Basal forebrain mediates motivational recruitment of attention by reward-associated cues. Frontiers in neuroscience 12:786.

Wietek J, Beltramo R, Scanziani M, Hegemann P, Oertner TG, Wiegert JS (2015) An improved chloride-conducting channelrhodopsin for light-induced inhibition of neuronal activity in vivo. Scientific reports 5:14807.

Woolf NJ, Hernit MC, Butcher LL (1986) Cholinergic and non-cholinergic projections from the rat basal forebrain revealed by combined choline acetyltransferase and Phaseolus vulgaris leucoagglutinin immunohistochemistry. Neuroscience letters 66:281–286.

Xu M, Chung S, Zhang S, Zhong P, Ma C, Chang W-C, Weissbourd B, Sakai N, Luo L, Nishino S (2015) Basal forebrain circuit for sleep-wake control. Nature neuroscience 18:1641–1647.

Zaborszky L, Carlsen J, Brashear HR, Heimer L (1986) Cholinergic and GABAergic afferents to the olfactory bulb in the rat with special emphasis on the projection neurons in the nucleus of the horizontal limb of the diagonal band. Journal of Comparative Neurology 243:488–509.

Zaborszky L, Duque A (2000) Local synaptic connections of basal forebrain neurons. Behavioural brain research 115:143–158.

Zhang K, Chen CD, Monosov IE (2019) Novelty, salience, and surprise timing are signaled by neurons in the basal forebrain. Current Biology 29:134–142.

Zheng Y, Feng S, Zhu X, Jiang W, Wen P, Ye F, Rao X, Jin S, He X, Xu F (2018) Different subgroups of cholinergic neurons in the basal forebrain are distinctly innervated by the olfactory regions and activated differentially in olfactory memory retrieval. Frontiers in neural circuits 12:99.

Zheng Y, Tao S, Liu Y, Liu J, Sun L, Zheng Y, Tian Y, Su P, Zhu X, Xu F (2022) Basal Forebrain-Dorsal Hippocampus Cholinergic Circuit Regulates Olfactory Associative Learning. International Journal of Molecular Sciences 23:8472.

